# The *Drosophila fussel* gene is required for bitter gustatory neuron differentiation acting within an Rpd3 dependent chromatin modifying complex

**DOI:** 10.1101/481721

**Authors:** Mathias Rass, Svenja Oestreich, Severin Guetter, Susanne Fischer, Stephan Schneuwly

## Abstract

Members of the Ski/Sno protein family are classified as proto-oncogenes and act as negative regulators of the TGF-ß/BMP-pathways in vertebrates and invertebrates. A newly identified member of this protein family is *fussel* (*fuss*), the *Drosophila* homologue of the human *functional Smad suppressing elements* (*fussel-15* and *fussel-18*). We and others have shown that Fuss interacts with SMAD4 and that overexpression leads to a strong inhibition of Dpp signaling. However, to be able to characterize the endogenous Fuss function in *Drosophila melanogaster,* we have generated a number of state of the art tools including anti-Fuss antibodies, specific *fuss*-Gal4 lines and *fuss* mutant fly lines via the CRISPR/Cas9 system. Fuss is a predominantly nuclear, postmitotic protein, mainly expressed in interneurons and *fuss* mutants are fully viable without any obvious developmental phenotype. To identify potential target genes or cells affected in *fuss* mutants, we conducted targeted DamID experiments in adult flies, which revealed the function of *fuss* in bitter gustatory neurons. We fully characterized *fuss* expression in the adult proboscis and by using food choice assays we were able to show that *fuss* mutants display defects in detecting bitter compounds. This correlated with a reduction of gustatory receptor gene expression (Gr33a, Gr66a, Gr93a) providing a molecular link to the behavioral phenotype. In addition, Fuss interacts with Rpd3, and downregulation of *rpd3* in gustatory neurons phenocopies the loss of Fuss expression. Surprisingly, there is no colocalization of Fuss with phosphorylated Mad in the larval central nervous system, excluding a direct involvement of Fuss in Dpp/BMP signaling.

Here we provide a first and exciting link of Fuss function in gustatory bitter neurons. Although gustatory receptors have been well characterized, little is known regarding the differentiation and maturation of gustatory neurons. This work therefore reveals Fuss as a pivotal element for the proper differentiation of bitter gustatory neurons acting within a chromatin modifying complex.

## Introduction

During development, the TGF-ß superfamily plays an important role in cell proliferation, differentiation, apoptosis, cell adhesion, wound healing, bone morphogenesis and cell motility [1]. Accordingly, there are multiple inhibitory factors taking care of proper regulation of TGF-ß pathways. Besides inhibitory Smads and Smurfs, which act by preventing the activation of TGF-ß receptors, another group of negative regulators of the TGF-ß pathway exists: The Ski/Sno protein family [2–4]. Although Ski/Sno proteins are classified as proto-oncogenes, their exact role in cancer progression is not fully understood. Various experimental approaches have identified pro- and anti-oncogenic features, where the tumor promoting function of Ski/Sno proteins seems to be mainly linked to their ability to counteract the anti-proliferative effects of TGF-ß signaling [5,6]. Physiologically, Ski and Sno have both been implicated in axonal morphogenesis, myogenesis and mammary gland alveogenesis [7–9]. Proteins of the Ski/Sno family are characterized by a Ski/Sno homology domain and a SMAD4 binding domain. These domains, although resembling DNA binding domains, mediate protein-protein interactions enabling binding of mSin3a, N-CoR, the histone deacetylase HDAC1, SMAD4 and different regulatory SMADs, thus leading to the recruitment of a repressive transcription complex, binding to target genes of the TGF-ß signaling pathway [10–12].

Whereas Ski and Sno are expressed mainly ubiquitously, two additional members of the Ski/Sno protein family, the functional smad suppressing elements (Fussel) 15 and 18 (Skor1 and Skor2 in mouse, respectively), are highly restricted to postmitotic neurons such as Purkinje cells [13–15]. Previous analysis showed that Skor1 interacts with Smad3 and acts as a transcriptional corepressor for LBX1, whereas Skor2 inhibits BMP signaling in overexpression assays and is required for the expression of Sonic Hedgehog in Purkinje cells. In addition, Skor2 is needed for proper differentiation of Purkinje cells and knockout mice die prematurely within 24 h after birth. However, there is no further insight into the functional mechanisms of Skor1 in mice [15–17].

In contrast to vertebrates, *Drosophila melanogaster* has only two Ski/Sno proteins: The Ski novel oncogene Snoo and the functional Smad suppressing element Fussel (Fuss). Snoo is the homologue of Ski and Sno and Fuss the homologue of Skor1 and Skor2. As in mice, Snoo is expressed broadly during development and adulthood, whereas Fuss expression is limited to a subset of cells in the nervous system during development [18]. Snoo has been found to be involved in eggshell patterning and in wing and tracheal development in *Drosophila melanogaster* [19–21]. Recent findings of Fuss function are highly controversial due to the lack of a reliable *Drosophila fussel* mutation. Loss of function experiments with a chromosomal deletion suggested, that Fuss acts as a cofactor for Smox signaling enabling ecdysone receptor (EcRB1) expression in developing mushroom bodies in the brain. In addition, a severe malformation of the adult mushroom bodies was detected [22]. Contrary to these results, in overexpression assays Fuss leads to an inhibition of BMP signaling via its interaction with Medea, the SMAD4 homologue in *Drosophila melanogaster* [18].

To clarify the molecular and physiological function of Fuss, we generated a complete loss of function allele via CRISPR/Cas9 editing. Interestingly, *fuss* mutants are fully viable, suggesting a modulatory function during development or/and adulthood, rather than an essential role for cell survival. To be able to better analyze Fuss expressing cells, we generated specific antibodies and reporter lines, which enabled us to further clarify the expression pattern of Fuss. During development, Fuss is expressed postmitotically in a highly restricted number of interneurons of the central nervous system (CNS). In our *fuss* mutant, we could show that Fuss, in contrast to previous studies, is neither acting as a negative regulator of BMP signaling endogenously, nor involved in mushroom body development. A targeted DamID (TaDa) experiment could not only confirm our findings molecularly, but also revealed, that Fuss is expressed in bitter sensory neurons. In consequence, *fuss* mutant flies lack the ability to sense bitter compounds and show reduced expression of bitter gustatory receptors. Furthermore, interaction studies show that Fuss can form a protein complex with Rpd3, a homologue of HDAC1, and indeed, downregulation of *rpd3* in bitter gustatory neurons resembles loss of Fuss. Thus, we propose that the Fuss/Rpd3 complex is required for proper cell fate determination in gustatory neurons either by direct or indirect control of the expression of gustatory bitter sensing receptors.

## Results

### Generation and analysis of *fuss* mutants

The *fuss* gene is localized on the fourth chromosome and therefore, due to the limited genetic resources for this chromosome, *fuss* mutations have escaped discovery in genetic screens for developmental mutations and previous research on this gene focused either on overexpression studies or on a chromosomal deletion covering multiple genes [18, 22]. However, it was shown recently that genes of the fourth chromosome can be CRISPR/Cas9 edited via homology directed repair and thus, we decided to generate a *fuss* null allele using this system [23]. A prerequisite for proper mutant generation is a detailed analysis of the genomic organisation of the gene. The *fuss* gene locus is fairly complex as it overlaps N-terminally with the RNA gene *sphinx* and C-terminally with the RNA gene *CR44030*. In addition, the Pax6 homologue *twin of eyeless* (*toy*) lies downstream of *fuss* and is transcribed in the opposite direction. Although the two genes are over 10 kb away from each other, regulatory sites of *toy* are located in the vicinity of the transcription start site of *fuss* [24].

Three different transcriptional start sites of *fuss* are annotated, which lead to three transcripts *fussB*, *fussC* and *fussD*. *FussB* and *fussD* have an identical amino acid sequence in contrast to the *fussC* transcript, which differs in 25 amino acids N-terminally from *fussB* and *fussD*. *FussC* uses an artificial promoter sequence due to a transposon insertion, leading to the assumption that this transcript is rather insignificant as it is of very low abundance (see below). To reduce side effects by deleting a silencer/enhancer structure of *sphinx* or *toy* and to maximise the effect on *fuss,* we removed an 855bp fragment, which is shared by all three transcripts (Fig 1A). This fragment contains the conserved Ski/Sno/Dac homology and SMAD4 binding domains, which are important for protein interactions and function of the Ski/Sno proteins [25,26]. Simultaneously, an attP site was introduced in the open reading frame of all *fuss* transcripts, which additionally results in a premature stop codon. Successful deletion of the two domains and the presence of the attP site was confirmed by PCR and subsequent DNA sequencing (S1A Fig). We termed this deletion *fuss*^*delDS*^ and it is a null allele. Due to the deletion and the premature stop codon no functional proteins can be made, which could be shown by anti-Fuss antibody stainings of heterozygous and homozygous *fuss*^*delDS*^ embryos (S1B Fig and S1C Fig). As a second mutant allele, we used the MiMIC-line *fuss*^*Mi13731*^, which is a gene trap insertion leading to the expression of GFP under the *fussB* and *fussD* promotor and a premature transcriptional stop of the *fussB* and *fussD* transcripts (Fig 1A). qPCR revealed that transcript levels of *fussB* and *fussD* in homozygous *fuss*^*Mi13731*^ flies are reduced to ten percent in contrast to WTB flies (S1D Fig). With an anti-GFP and anti-Fuss antibody staining, we could confirm that heterozygous *fuss*^*Mi13731*^*/+* flies express GFP in the correct Fuss expression pattern, whereas Fuss staining in homozygous *fuss*^*Mi13731*^ flies is reduced to background levels (S1E Fig and S1F Fig). We conclude that *fuss*^*Mi13731*^ is at least a strong hypomorph for *fuss*, which further suggests that *fussC* is not specifically expressed or only at very low levels. In addition to *fuss*^*Mi13731*^, which can be used as a GFP reporter line, we created a Gal4 line from *fuss*^*Mi13731*^ by recombination mediated cassette exchange [27]. This Gal4 line was named *fussBD*-Gal4 and it was edited with the CRISPR/Cas9 system following the same strategy as for the *fuss*^*delDS*^ allele. This resulted in a line called *fuss*^*delDS*^-Gal4, which allows Gal4 expression in Fuss expressing cells in a mutant background enabling us to analyze the presence and integrity of these cells. At last, we generated a UAS*-t::gRNA-fuss*^*4x*^, which allows cell specific gene disruption of *fuss* via the UAS-Gal4 system [28]. The gRNAs target four CRISPR target sites located in the DNA sequence of the Ski/Sno homology domain (S1G Fig). We could detect a strong loss of GFP signal in adult brains of flies overexpressing GFP tagged Fuss, Cas9 and *t::gRNA-fuss*^*4x*^ by *fussBD*-Gal4 compared to adult brains only expressing GFP tagged Fuss and Cas9 by *fussBD*-Gal4 (S1H and S1I Fig).

**Fig 1.**
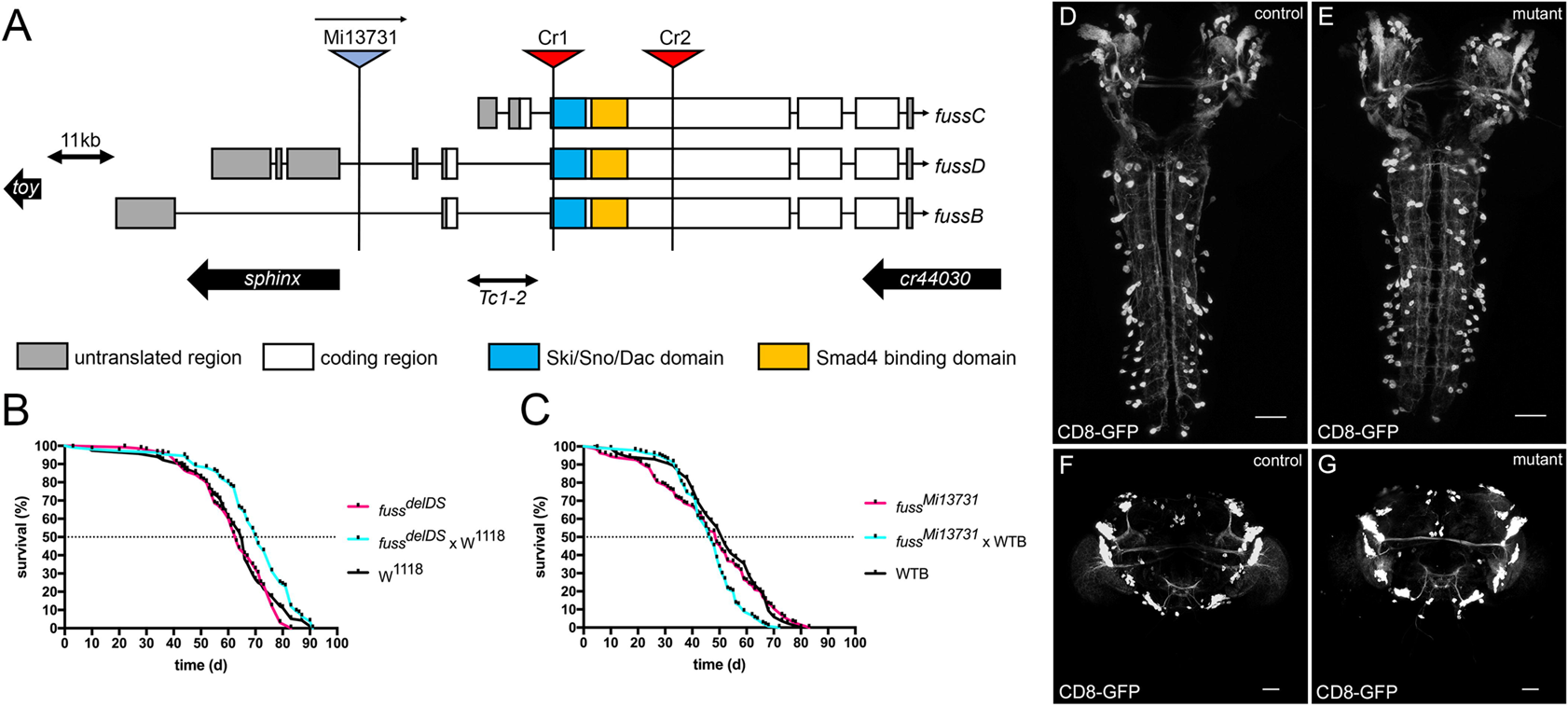
Generation of CRISPR/Cas9 induced *fuss* mutant. (A) Generation of a *fuss* knockout allele, which lacks the Ski/Sno/Dac homology domain (blue box) and the SMAD4 binding domain (yellow box) by means of two CRISPR target sites Cr1 and Cr2. The location of the second mutant allele *fuss*^*Mi13731*^, a gene trap insertion in the FussB and FussD transcript is indicated. Approximate locations of *toy*, *sphinx* and *cr44030* are depicted as black arrows. The location of the transposable element *Tc1-2* is shown as a double-sided arrow. (B) Differences in the mean lifespan of homozygous *fuss*^*delDS*^ flies (n=133, purple) and the two controls W^1118^ (n=116, black) and *fuss*^*delDS*^ x W^1118^ flies (n=109, blue) are not relevant. C) Longevity experiments show no differences between homozygous *fuss*^*Mi13731*^ (n=164, purple), heterozygous *fuss*^*Mi13731*^ x WTB (n=136, blue) and WTB flies (n=85, black). (D) Projection pattern and cell bodies of larval brains visualized by expression of UAS-*CD8-GFP* by *fuss*^*delDS*^-Gal4/*+* larvae (control). (E) Projection pattern and cell bodies of larval brains visualized by expression of UAS-*CD8-GFP* by *fuss* mutant *fuss*^*delDS*^-Gal4/*fuss*^*delDS*^ larvae (mutant). (F) Projection pattern and cell bodies of adult brains visualized by expression of UAS-*CD8-GFP* by *fuss*^*delDS*^-Gal4/+ flies (control). (G) Projection pattern and cell bodies of adult brains visualized by expression of UAS-*CD8-GFP* by *fuss* mutant *fuss*^*delDS*^-Gal4/*fuss*^*delDS*^ flies (mutant). Scale bars indicate 50 *µ*m.

In a previous study by Takaesu et al. [22], a 40 kb spanning genomic deletion including the *fuss* gene (among several other genes) was used for functional studies of the *fuss* gene. They observed a strongly reduced survivability during development and a decreased lifespan, which was attributed to the loss of Fuss expression alone. In contrast to their results, we did not observe a reduced survivability during larval or pupal stages with our *fuss* mutant flies. Therefore, we conducted longevity experiments. Neither homozygous *fuss*^*Mi13731*^ nor *fuss*^*delDS*^–flies showed a significant reduction in lifespan compared to their controls (Fig 1B and Fig 1C).

Next, we compared the CD8-GFP expression pattern of heterozygous *fuss*^*delDS*^-Gal4/+ and mutant *fuss*^*delDS*^-Gal4/*fuss*^*delDS*^ flies. We did neither observe an evident loss of GFP positive cells in the CNS of third instar larvae (Fig 1D and Fig 1E) nor in three to five-day old adult flies (Fig 1F and Fig 1G). Therefore, loss of fuss does neither lead to cell death, nor to a reduced survival during development or to a shortened lifespan.

### Characterization of embryonic Fuss expressing cells reveals distinct neuronal identities

Due to the absence of any clear visible phenotype, we created specific polyclonal antibodies against a 16 kDa nonconserved fragment localized at the C-terminus of Fuss to characterize Fuss expressing cells and to draw conclusions about its function (S2A Fig). These anti-Fuss antibodies clearly detect a Fuss-GFP fusion protein on western blots and stainings mirror previously conducted RNA *in situ* hybridisations (S2B Fig) [18, 22].

In a first overview of Fuss staining during embryonic development, Fuss expression is mainly observed in the embryonic brain (Fig 2A, circles), the developing stomatogastric nervous system (Fig 2A, arrowhead), single cells lying anterior to the CNS (Fig 2A arrows), which will develop to inner gustatory neurons as shown later and the ventral nerve cord (VNC, Fig 2B). As Fuss is characterized by its conserved domains as a member of the Ski/Sno protein family, which are all considered to be transcription regulators, we observe Fuss protein, as expected, exclusively localized in the nucleus. During embryonic development, Fuss protein appears first at stage 13 and the number of Fuss positive cells increases continuously from early to late embryonic stages as previously observed (S2C Fig [22]). At embryonic stage 16, expression can be observed in two to five cells per hemineuromer with ascending numbers from posterior to anterior (Fig 2B and S2C Fig).

**Fig 2.**
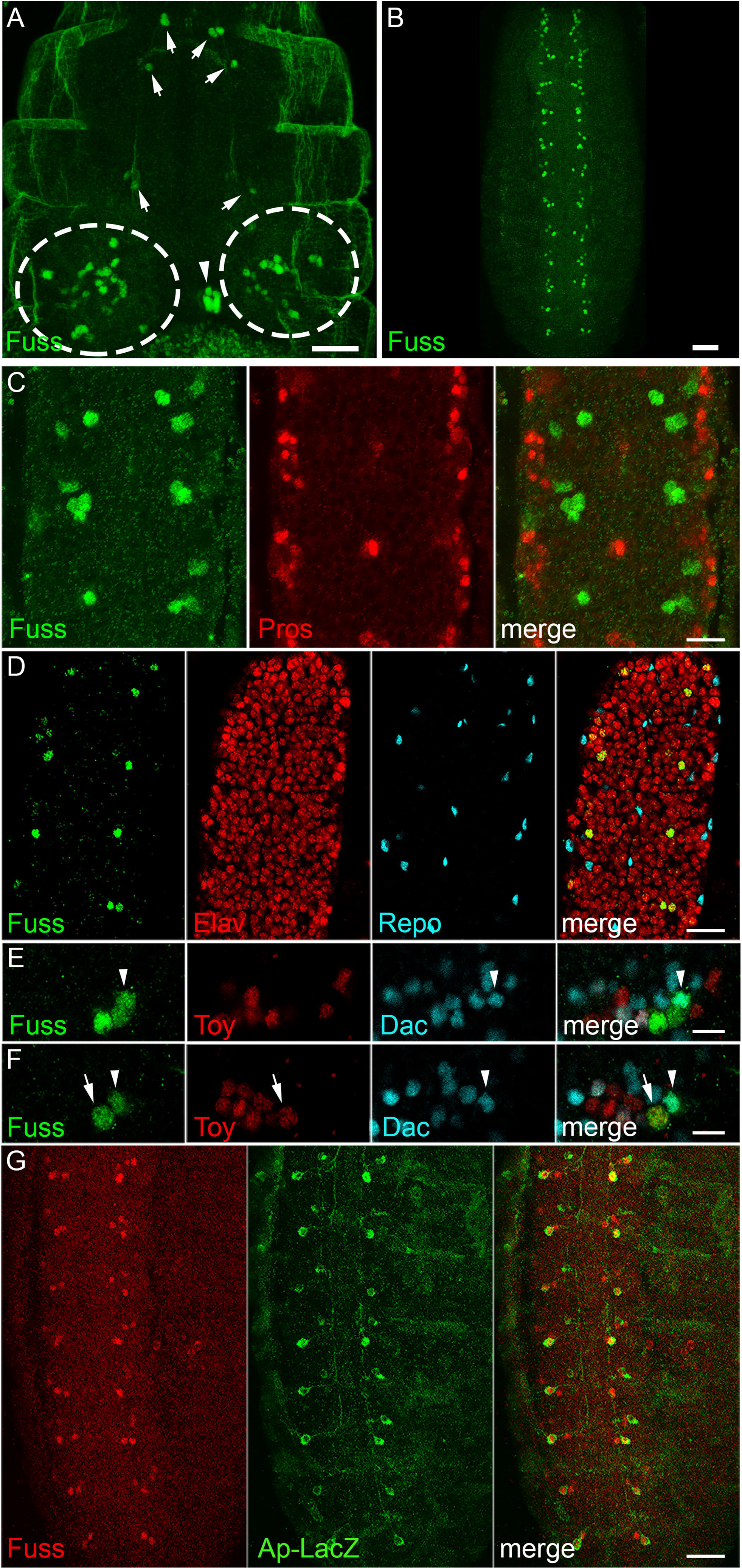
Fuss is expressed in postmitotic interneurons. (A) In stage 16 embryos, anti-Fuss staining can be observed in a restricted number of cells in the CNS (dashed circles). Fuss is also found in individual cells of the stomatogastric nervous system (arrowhead) and anterior to the CNS (arrows). (B) In the ventral nerve cord (VNC), Fuss is expressed in two to five neurons per hemineuromer in ascending number from posterior to anterior. (C) Confocal microscopy images reveal that Fuss expressing cells in the VNC (green) do not overlap with ganglion mother cells stained with anti-Prospero (red). (D) Fuss is exclusively expressed in neurons (Elav, red) but not in glia (Repo, blue). (E, F) In one representative hemineuromer five Fuss (green) positive neurons are characterized regarding their Dac (blue, arrowhead) or Toy (red, arrow) expression. (G) In every hemineuromer one Fuss (green) neuron is positive for LacZ (red) expressed under the *apterous* promotor. Scale bars indicate 25*µ*m (A), 50*µ*m (B), 10 *µ*m (C, E, F) and 20 *µ*m (D, G).

The late appearance of the Fuss protein during development suggested, that Fuss might be expressed only postmitotically. We confirmed this hypothesis by visualizing ganglion mother cells in the embryo with anti-Prospero and Fuss cells with anti-Fuss antibodies and no overlapping stainings were detected (Fig 2C). As shown by colocalization studies with the glia marker Repo and the neuronal marker Elav, the staining pattern is exclusively neuronal (Fig 2D). To further identify neuronal subpopulations in hemineuromers of the VNC, prominent neuronal markers such as Engrailed (En), Even skipped (Eve), Apterous (Ap), Hb9, Dachshund (Dac) and Twin of eyeless (Toy) were utilised. No colocalization of Fuss with the interneuron marker En or with the motoneuron markers Eve or Hb9 was observed (S2D Fig, S2E Fig and S2F Fig). Because Eve and Hb9 label most of the embryonic motoneurons, Fuss is unlikely to be expressed in motoneurons [29,30]. We were especially interested if Dac and Fuss colocalize, because the interneuron marker Dac shares sequence similarity with Ski and Sno and consequently is a related protein to Fuss [31,32]. Interestingly, Dac and Fuss are partially coexpressed, which emphasizes that at least some Fuss neurons are interneurons (arrowhead, Fig 2E and Fig 2F). As the Toy gene lies only 11 kb downstream of Fuss as it is transcribed in the opposite direction, it is reasonable that they partially share enhancer/silencer regions. Remarkably, we only found one Toy positive Fuss neuron per hemineuromer in the VNC (arrow, Fig 2E and Fig 2F) excluding extensive overlap of regulatory regions. Ap is expressed in three cells per abdominal hemineuromer. These cells are subdivided into one dorsal Ap and two ventral Ap interneurons [33]. Using the *ap*-tau-LacZ reporter, which only labels one ventral Ap cell and the *ap*-Gal4 driver line we showed, that both ventral AP interneurons are Fuss positive (Fig 2G and S2G Fig). Due to the location of the Toy positive Fuss neuron, we assume that it is one of the ventral Ap cells and therefore also an interneuron.

Taken together we could show that Fuss is expressed only postmitotically in interneurons in the developing CNS, which will be further confirmed later.

### Transcriptional profiling of adult Fuss neurons in the head

Fuss is expressed in heterogenic neuronal populations, which are represented by differentially expressed markers and by their projection patterns. To develop new approaches to identify and study viable phenotypes in *fuss* mutants, it was of upmost importance to identify genes, which are regulated by Fuss. Therefore, we performed a targeted DamID (TaDa) experiment by expressing a Dam-PolII fusion protein with the *fuss*^*delDS*^-Gal4 driver line. RplI215, the large subunit of the RNA Polymerase II, is fused with the Dam methylase and thus this so called Dam-PolII fusion protein enabled us to detect the binding sites of the RNA Polymerase II similar to an RNA PolII ChIP and to detect transcribed genes in these neurons without cell sorting [34]. As a control, the unfused Dam protein was expressed with the *fuss*^*delDS*^-Gal4 driver line. Expression of UAS-*Dam* or UAS-*Dam-PolII* was inhibited by Gal80ts during development and expression of this proteins was allowed for 24 h at 29 °C in one to three-day old flies. Next generation sequencing libraries were generated from three different biological replicates expressing Dam-PolII and from three replicates expressing Dam alone. Each experiment was compared to each control leading to nine individual datasets. Because the binding patterns of all nine files were highly similar, individual datasets were averaged to reduce the amount of false positive hits of expressed genes. Genes with a false discovery rate (FDR) lower than 0.01 were accounted as expressed resulting in 2932 genes (S1 Appendix). The TaDa data is represented as a log2 ratio of Dam-PolII/Dam. As expected, *fuss* was one of the genes with the lowest FDR and highest PolII coverage (Fig 3A). This clearly indicates that the approach was carried out successfully. Furthermore, genes already identified by antibody stainings such as *elav*, *dac* or *toy*, were also detected by the TaDa experiment. Toy was also expressed in some Fuss neurons in adult brains (Fig 3B). This again underlines, like already observed during embryonic development, that *fuss* and *toy* might share common silencer/enhancer elements with Fuss. To further verify the TaDa data, colocalization experiments were conducted. Two cell fate markers *atonal* (*ato*) and *acj6* were enriched in our dataset and we could also detect the expression of these two proteins via immunofluorescense stainings in Fuss neurons (Fig 3B). Furthermore, we analyzed *genes* which show no or low PolII coverage e.g. *pale* (*ple*) and *Insulin-like peptide 2* (*Ilp2*) via immunofluorescence and could not detect any staining in Fuss positive neurons (S3 Fig). In particular, the absence of Fuss in insulin producing cells is in disagreement with recent published results using enhancer/reporter constructs (S3B Fig, [35,36]). In summary, we can conclude, that using this strategy, we have successfully generated an adult Fuss neuron specific transcriptional profile.

**Fig 3.**
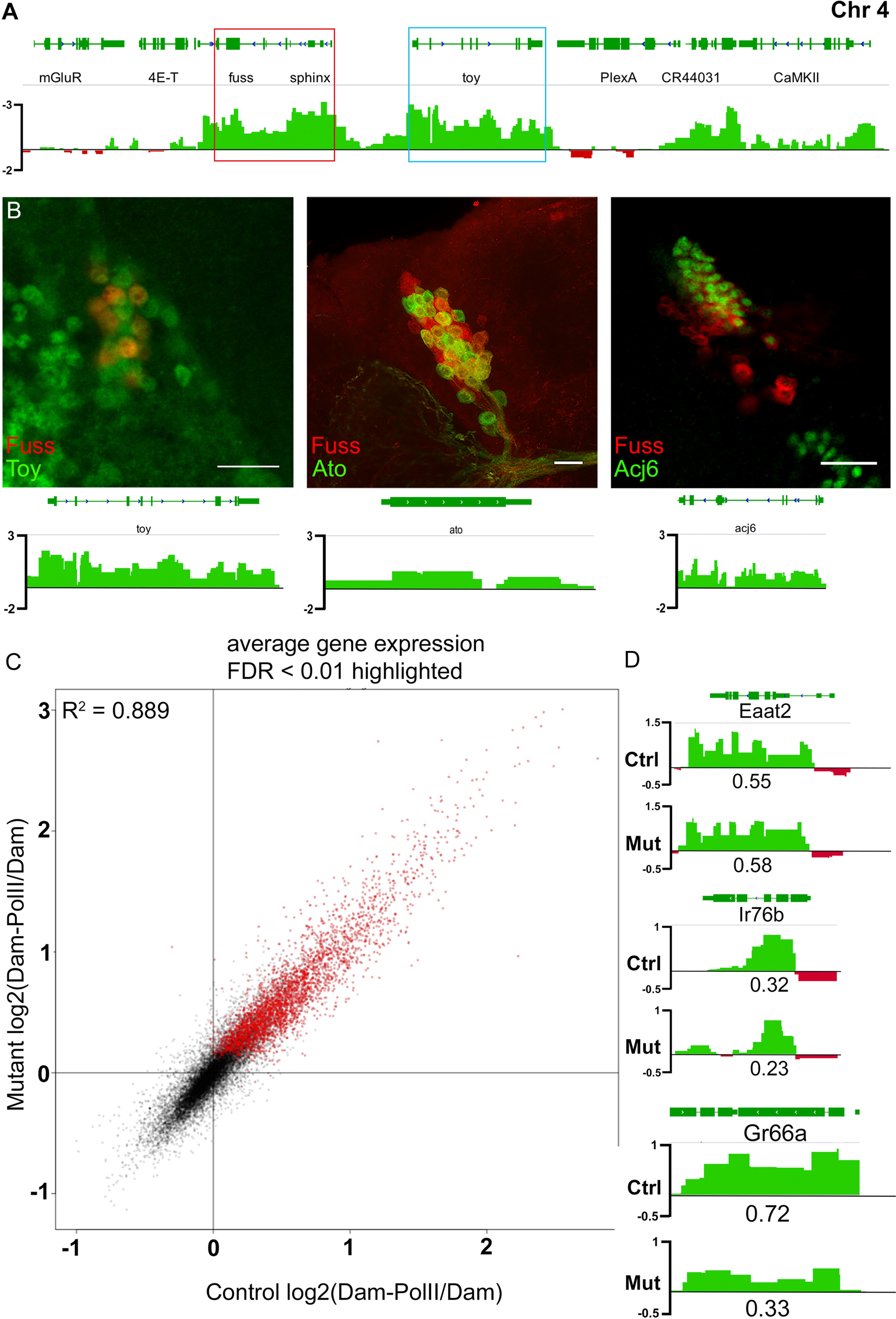
Targeted DamID of control and *fuss* mutants reveal sensory neuron markers as potential Fuss targets. (A) A Dam-PolII/Dam binding pattern was generated from nine individual TaDa profiles and averaged to one single track. *fuss* (red box) and *toy* (blue box) are highly covered by Dam-PolII. Regions bound stronger by Dam-PolII than by Dam are depicted in green. Regions bound stronger by Dam than by Dam-PolII are depicted in red. (B) Verification of three TaDa positive genes, *toy*, *atonal* and *acj6* by immunostaining. Toy was labelled by anti-Toy (green) and Fuss by anti-Fuss staining (red). Expression of LacZ by *ato*-Gal4 (green) and expression of GFP from the *fuss*^*Mi13731*^/+ reporter line (red). Labelling of Acj6 with anti-Acj6 antibody (green) and expression of CD8-GFP with *fussBD*-Gal4 (red). Scale bars indicate 10 *µ*m. (C) log2(Dam-PolII/Dam) data from controls compared with log2(Dam-PolII/Dam) data from mutant Fuss neurons show only small deviations from each other. Coefficient of determination R^2^ = 0.889. (D) TaDa reveals sensory neuron marker expression of *EAAT2*, *Ir76a*, and *Gr66a* in both datasets (upper lane: control; lower lane: mutant) with a clear reduction in PolII coverage of *Gr66a* in the mutant dataset in contrast to the control dataset. PolII coverage is depicted under PolII binding pattern. Regions bound stronger by Dam-PolII than by Dam are depicted in green. Regions bound stronger by Dam than by Dam-PolII are depicted in red.

In the next step, we wanted to search for potential target genes of Fuss using the same strategy and conditions as above, but this time *fuss*^*delDS*^-Gal4 was kept over the *fuss*^*delDS*^ allele to profile transcription of *fuss* mutant neurons. Again, individual datasets were averaged and genes with an FDR lower than 0.01 were accounted as expressed resulting in 3150 genes (S2 Appendix). The comparison of the log_2_(DamPolII/Dam) data of heterozygous *fuss*^*delDS*^/+ and homozygous *fuss*^*delDS*^ flies showed, that there is not a strong deviation (coefficient of deviation R^2^ = 0.889) of the mutant transcriptional profile from the control (Fig 3C). Because Fuss is only expressed in a small number of CNS neurons, the acquired data can only be confirmed by antibody staining and not by semiquantitive qPCR or western blots from whole heads. There were three genes which attracted our attention: *Eaat2*, *Ir76b* and especially *Gr66a* as they provided a possible link to Fuss expression in gustatory sense neurons (Fig 3D). These genes could be found in both datasets, although only *Eaat2* had an FDR lower than 0.01 in both datasets. The PolII coverage of *Eaat2* and *Ir76b* was only slightly different between homozygous and heterozygous flies, whereas *Gr66a*, which is exclusively expressed in bitter gustatory sense neurons (GRNs), showed a significant reduction in mutant flies (S1 Appendix and S2 Appendix).

### Fuss is expressed in a subset of gustatory neurons

It has been shown that the glutamate aspartate transporter Eaat2 is expressed in sensory neurons [37]. The ionotropic receptor Ir76b is expressed in gustatory neurons and the gustatory receptor Gr66a is specifically expressed in bitter GRNs, where Gr66a is a very important component for bitter taste sensation [38,39]. We already observed Fuss expression in cells outside of the larval CNS, therefore, to confirm the TaDa datasets, we analyzed gustatory neurons in larval and adult stages. In larvae, Fuss expression cannot be observed in the terminal or dorsal organ, but it can be found in the inner gustatory sense organs. We found Fuss expression in two pairs of neurons in the dorsal pharyngeal sensilia (DPS, Fig 4A) one neuron pair in the dorsal pharyngeal organ (DPO, Fig 4A) and two neuron pairs in the posterior pharyngeal sensilia (PPS, Fig 4A). None of the GRNs in the ventral pharyngeal sense organ (VPS) express Fuss. These cells have been already characterized by expression of different gustatory receptors and we found that larval Fuss expressing GRNs show a colocalization with a marker for bitter sensing neurons Gr33a [40]. In addition, one neuron pair in the DPS also shows an overlap with Gr93a which has been shown to be important for caffeine response in larvae (Fig 4B, [41,42]).

**Fig 4.**
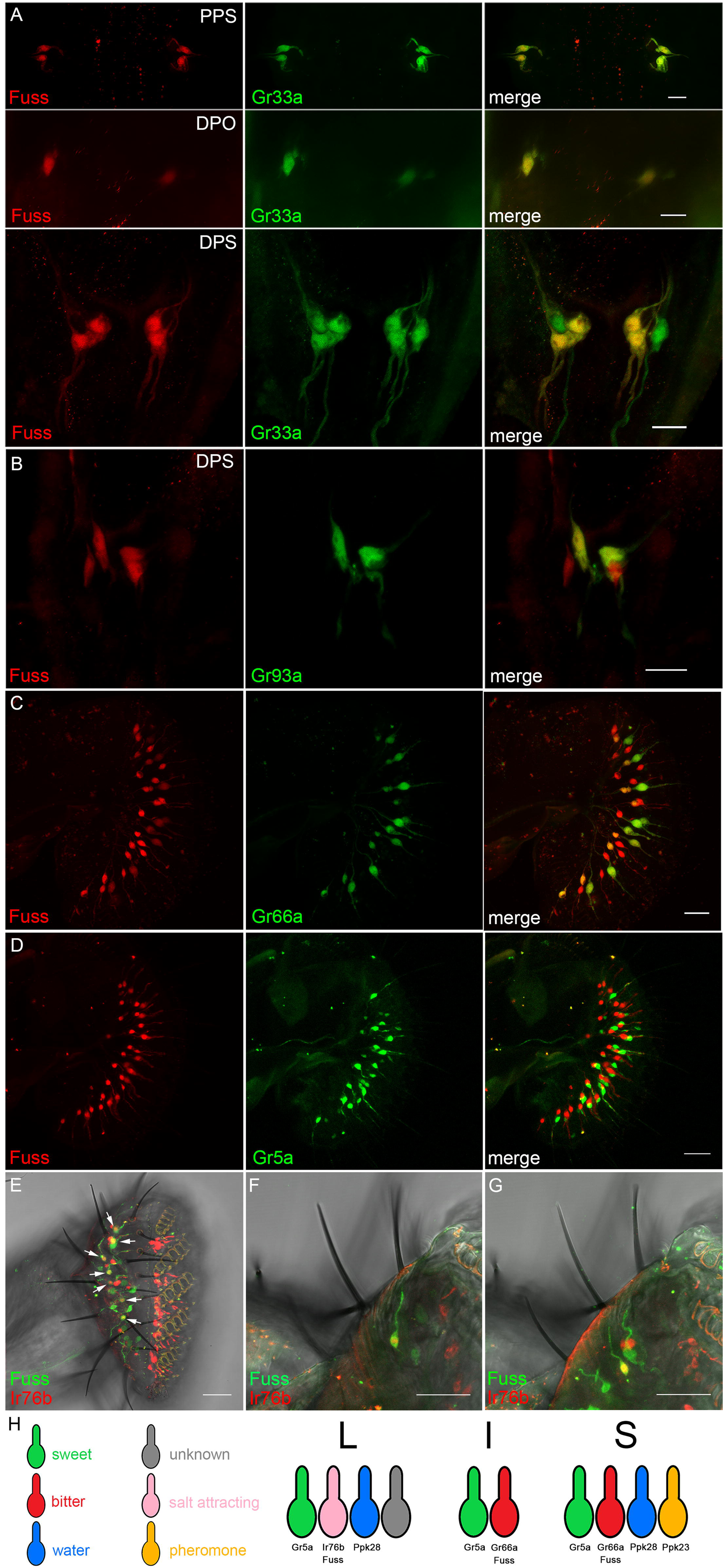
Fuss is expressed in bitter GRNs in inner gustatory organs of larvae and in bitter and salt attracting GRNs of the adult proboscis. (A) GFP (red) expression from heterozygous *fuss*^*Mi13731*^/+ reporter line can be observed in bitter gustatory neurons marked with LacZ (green) expressed via *Gr33a*-Gal4 in the dorsal (DPS), posterior pharyngeal sensilia (PPS) and dorsal pharyngeal organ (DPO) identifying Fuss neurons as bitter gustatory neurons in these organs. (B) LacZ (green) expressed from a *Gr93a*-Gal4 line colocalizes with GFP (red) from a heterozygous *fuss*^*Mi13731*^/+ reporter line. Scale bars indicate 10 *µ*m. (C) In adult flies, GFP (red) expression from the *fuss*^*Mi13731*^ gene trap line can be observed in the proboscis in one GRN per bristle. LacZ (green) driven by the bitter gustatory driver line *Gr66a*-Gal4 can be found in S- and I-type sensilla of GFP expressing GRNs. Scale bar indicates 20 *µ*m. (D) No overlap between LacZ (green) driven by Gr5a-Gal4 and GFP (red) expressed from *fuss*^*Mi13731*^/+ can be observed. (E-G) LacZ (red) expressed by *Ir76b*-Gal4 and GFP (green) expressed from *fuss*^*Mi13731*^/+ overlap in L-type sensilla (arrows). Scale bars indicate 20 *µ*m (C, D) and 25 *µ*m (E-G), respectively. (H) Schematic representation of *fuss* expression in GRNs of L-, I- and S-type sensilla.

Later, in adulthood, Fuss expression continues in GRNs of the proboscis. In the adult labellum three different types of sensilla can be found divided into short (S-type), intermediate (I-type) and long sensilla (L-type). Intermediate sensilla are innervated by two GRNs and short and long sensilla by four GRNs [43]. Interestingly Fuss expression is observed in one GRN per gustatory sensilla and is consistently colocalized with the bitter GRN marker Gr66a in neurons innervating short and intermediate sensilia (Fig 4C [38]). Long sensilla do not contain a Gr66a positive GRN, therefore, all Gr66a neurons in the labellum are Fuss positive, but not vice versa. Another gustatory receptor which is broadly expressed and labels sweet GRNs is Gr5a, but no overlap with Fuss positive neurons was observed (Fig 4D). Besides Gr66a our TaDa dataset revealed that the ionotropic receptor Ir76b is expressed in Fuss neurons. Ir76b has been shown to be expressed by one GRN per L-type sensillum, which plays a role in attractive salt tasting [44]. We found that in L-type sensilla Fuss is coexpressed with Ir76b (Fig 4E-G). Besides the expression in GRNs of the proboscis we found Fuss being expressed in two GRNS each in the last two tarsal segments in every leg (S4A Fig). In conclusion, we integrated Fuss expression into the GRN model from Freeman and Dahanukar (Fig 4H, [45]) and demonstrate that Fuss is expressed in bitter neurons in S- and I-type sensilla and in salt attracting neurons in L-type sensilla.

### Loss of Fuss impairs bitter taste sensation

By its gustatory system *Drosophila melanogaster* can discriminate between valuable food sources for foraging or egg laying and toxic compounds which could harm the fly or its offspring [46]. To address if Fuss is required for the proper development of GRNs, we focused on the impact of Fuss mutation on differentiation of bitter GRNs, because Fuss is expressed in all bitter GRNs of the proboscis. To detect if *fuss* mutant flies display an impaired bitter taste sensation, we tested one to three-day old flies in a two-choice feeding assay. In our standard test, flies had to choose between 1mM sucrose or 5mM sucrose plus 10mM caffeine. We calculated a preference index ranging from zero to one, where zero indicates complete avoidance of the bitter compound and one a complete preference for it, due to the higher sugar concentration. First, Fuss expressing neurons were ablated by UAS-*rpr* expression with *fussBD*-Gal4 to show their importance in bitter sensing and indeed, these flies showed a strong impairment of bitter discrimination (Fig 5A). Furthermore, homozygous *fuss*^*Mi13731*^, *fuss*^*delDS*^ and transheterozygous mutants (*fuss*^*Mi13731*^/*fuss*^*delDS*^) as well as their appropriate controls were tested. All mutant genotypes showed an increased preference for 5mM sucrose mixed with caffeine and by overexpression of Fuss in *fuss* mutant neurons we could revert preference to wildtype levels (Fig 5A). To show that the behavioural phenotype of *fuss* mutants is due to defects in GRNs and not derived from other higher order Fuss neurons in the CNS we specifically disrupted *fuss* in all GRNs with the *Poxn*-Gal4-13-1 driverline and our UAS-*cas9*; UAS-*t::gRNA-fuss*^*4x*^ flies. *Poxn*-Gal4-13-1 expresses Gal4 early in development in all GRNs and in ellipsoid body neurons as well as interneurons of the antennal lobe of the brain (Fig S4B, [47]), therefore the only common neuronal populations between Fuss and *Poxn*-Gal4-13-1 are the GRNs and indeed, as shown in Fig 5A, these flies show the same bitter sensing deficits. We also tested different concentrations of caffeine as well as another bitter compound (denatonium benzoate) and *fuss*^*Mi13731*^ flies always displayed a higher preference towards the 5mM sucrose mixed with the bitter compound than controls except when concentration of the bitter compound was too high (S4C and S4D Fig). Thus, not only detection of caffeine but more general bitter sensation is disturbed, because different GR multimers are needed for the detection of different aversive compounds, e.g. Gr93a which is expressed in a subset of S-type sensilla is needed for caffeine but not for denatonium benzoate sensation [48]. The gustatory receptor *Gr66a* showed a strong reduction in PolII coverage in mutant flies in contrast to control flies and is only expressed in a proportion of Fuss positive GRNs. The gustatory receptor GR33a has been found to be coexpressed with Gr66a in bitter GRNs and both are involved in bitter sensation, particularly together with Gr93a in caffeine sensation [40,48]. To validate GRN results from the TaDa experiment, we extracted RNA from adult proboscis and analysed the expression levels of those GRs via semiquantitative RT-PCR. In homozygous *fuss*^*Mi13731*^-flies Gr33a and Gr66a expression were strongly reduced as compared to WTB and heterozygous *fuss*^*Mi13731*^-flies. Gr93a expression levels of homozygous *fuss*^*Mi13731*^-flies were similar to WTB levels but reduced when compared to heterozygous *fuss*^*Mi13731*^-flies (Fig. 5B). The observed effects were enhanced in *fuss*^*delDS*^-flies. Gr33a, Gr66a and Gr93a expression levels were all reduced in *fuss*^*delDS*^-flies in contrast to both controls (Fig. 5C). A similar downregulation of Gr33a, Gr66a and Gr93a expression levels was observed in transheterozygous *fuss*^*Mi13731*^/*fuss*^*delDS*^ flies in contrast to WTB flies (S4E Fig). Next, we tested if the number of Gr33a and Gr66a positive GRNs is reduced in *fuss* mutant flies. We counted Fuss positive and Gr33a positive neurons in flies of the genotypes *Gr33a*-Gal4/UAS-*LacZ*; *fuss*^*Mi13731*^/+ and *Gr33a*-Gal4/UAS-*LacZ*; *fuss*^*Mi13731*^/*fuss*^*delDS*^. In this genetic combination we counted 2.5 less Fuss positive cells and surprisingly 7.2 less Gr33a positive cells in controls than in transheterozygous mutants (Fig 5D). Furthermore, we analysed number of Fuss positive and Gr66a positive neurons in flies of the genotypes UAS-*LacZ*/+; *Gr66a*-Gal4/+; *fuss*^*Mi13731*^/+ and UAS-*LacZ*/+; *Gr66a*-Gal4/+; *fuss*^*Mi13731*^/*fuss*^*delDS*^. We found the same reduction in overall number of Fuss positive GRNs. But the number of Gr66a positive GRNs is decreased at the same level as the number of overall Fuss positive GRNs (Fig 5E). Thus, the overall number of bitter GRNs is slighty reduced in *fuss* mutant flies, but interestingly Gr33a expression is completely abolished in some bitter GRNs, whereas the reduction of Gr66a expression found in qPCR experiments does not result in a reduced number of Gr66a positive GRNs. So, upon the loss of Fuss expression, bitter GRN differentiation is highly disturbed, which renders these flies inable to detect bitter compounds.

**Fig 5.**
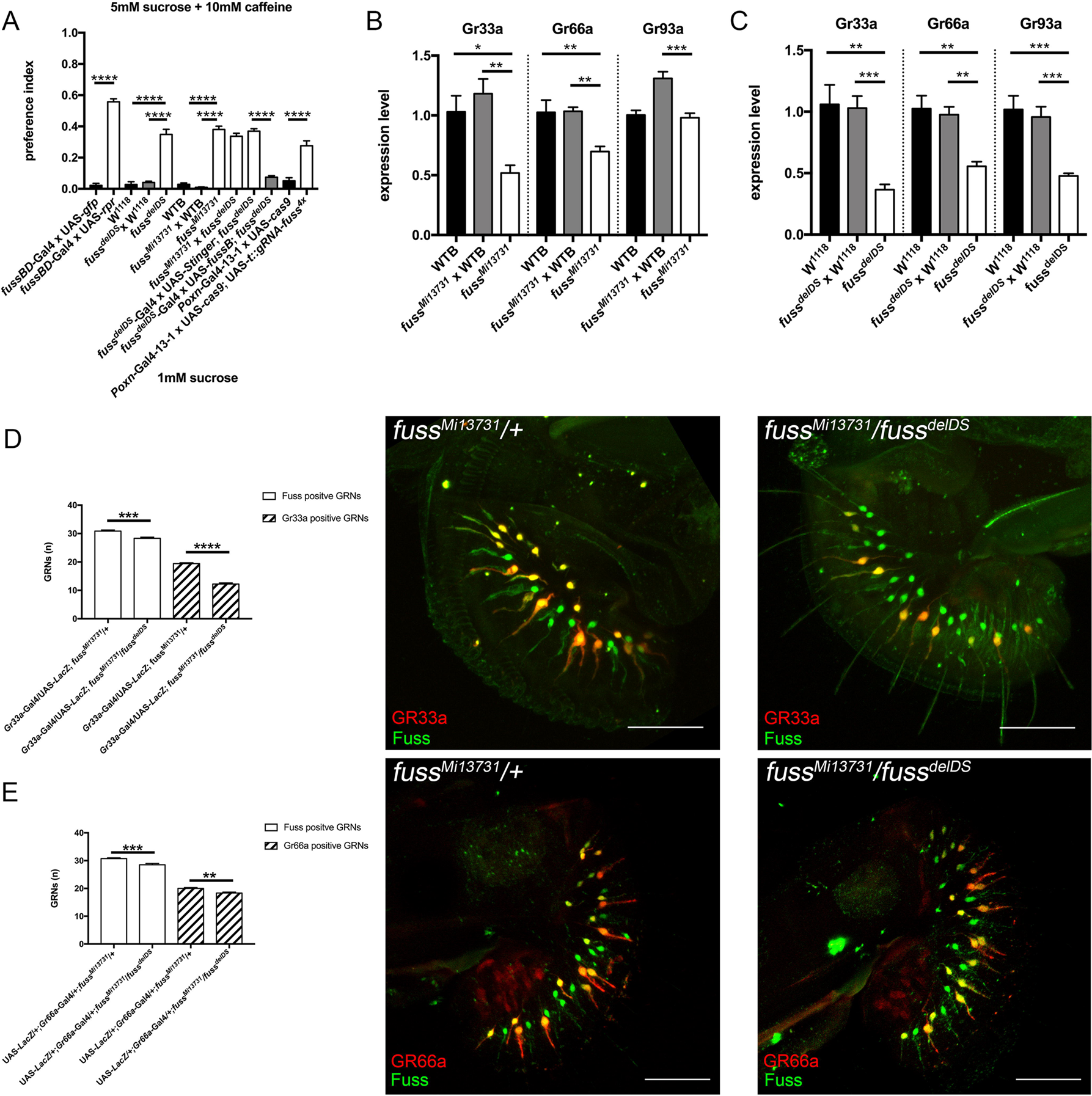
*fuss* mutant GRNs show impaired caffeine avoidance. (A) Two-choice feeding assay reveals reduced caffeine sensation of homozygous *fuss*^*Mi13731*^ and *fuss*^*delDS*^ flies in contrast to their appropriate controls. As a positive control, Fuss neurons were ablated by expression of *rpr* via *fussBD*-Gal4. Transheterozygous fuss^delDS^/fuss^Mi13731^ flies also have a reduced ability to sense bitter compounds comparable to levels of homozygous *fuss*^*Mi13731*^ and *fuss*^*delDS*^ flies. Overexpression of *fussB* with *fuss*^*delDS*^-Gal4 reduces caffeine preference to wildtype levels. Flies with a GRN specific *fuss* gene disruption (*Poxn*-Gal4-13-1 x UAS-*cas9*; UAS-*t::gRNA-fuss4x*) show a reduced caffeine sensation compared to controls (*Poxn*-Gal4-13-1 x UAS-*cas9*) (n=4-10 for each genotype). One-way ANOVA with *post hoc* Tukey’s test was used to calculate p-values. ****p<0.0001. Error bars indicate SEM. (B) Semiquantitative qPCR of bitter gustatory receptors Gr33a, Gr66a and Gr93a reveals a reduced expression of Gr33a and Gr66a in homozygous *fuss*^*Mi13731*^ flies in contrast to controls. Gr93a expression of homozygous *fuss*^*Mi13731*^ flies is only reduced if compared to heterozygous *fuss*^*Mi13731*^ x WTB but not WTB flies. n=4-6 for each genotype. One-way ANOVA with *post hoc* Tukey’s test was used to calculate p-values. *p<0.05. **p<0.01. ***p<0.001. Error bars indicate SEM. (C) Analysis of Gr33a, Gr66a and Gr93a expression by semiquantitative qPCR reveals reduced expression of GRs in homozygous *fuss*^*delDS*^ flies in contrast to heterozygous *fuss*^*delDS*^ x W^1118^ and W^1118^ flies (n=4-6 for each genotype). One-way ANOVA with *post hoc* Tukey’s test was used to calculate p-values. *p<0.05 **p<0.01 ***p<0.001. Error bars indicate SEM. (D) Comparison of Fuss positive neurons and Gr33a positive neurons of the genotypes *Gr33a*-Gal4/UAS-*LacZ*;*fuss*^*Mi13731*^/+ (control) and *Gr33a*-Gal4/UAS-*LacZ*;*fuss*^*Mi13731*^/*fuss*^*delDS*^ (mutant) shows a slight reduction in Fuss positive GRN numbers (30.8 vs 28.3) and a strong reduction in Gr33a positive GRN numbers (19.4 vs 12.2). Unpaired t-test was used to calculate p-values. n=12-13 for each genotype. ***p<0.001. ****p<0.0001. Error bars indicate SEM. Adult proboscis of genotypes *Gr33a*-Gal4/UAS-*LacZ*;*fuss*^*Mi13731*^/+ (control, abbr: *fuss*^*Mi13731*^/+) and Gr33a-Gal4/UAS-LacZ;*fuss*^*Mi13731*^/*fuss*^*delDS*^ (mutant, abbr: *fuss*^*Mi13731*^/ *fuss*^*delDS*^). Scale bar indicates 50 *µ*m. (E) Comparison of Fuss positive neurons and Gr66a positive neurons of the genotypes UAS-*LacZ*/+;*Gr66a*-Gal4/+;*fuss*^*Mi13731*^/+ (control) and UAS-*LacZ*/+;*Gr66a*-Gal4/+;*fuss*^*Mi13731*^/*fuss*^*delDS*^ (mutant) shows a slight reduction in Fuss positive GRN numbers (30.75 vs 28.5) and an equal reduction in Gr66a positive GRN numbers (20 vs 18.3). n=12 for each genotype. Unpaired t-test was used to calculate p-values. **p<0.01. ***p<0.001. Error bars indicate SEM. Adult proboscis of genotypes UAS-*LacZ*/+;*Gr66a*-Gal4/+;*fuss*^*Mi13731*^/+ (control, abbr: *fuss*^*Mi13731*^/+) and UAS-*LacZ*/+;*Gr66a*-Gal4/+; *fuss*^*Mi13731*^/*fuss*^*delDS*^ (mutant, abbr: *fuss*^*Mi13731*^/ *fuss*^*delDS*^). Scale bar indicates 50 *µ*m.

### Fuss interacts with the histone deacetylase Rpd3 to affect cell fate determination

In mammals there are two homologues of Fuss, Skor1 and Skor2, which display a high sequence conservation within the Ski/Sno/Dac homology domain and the SMAD4 binding domain. In contrast, the conservation in the C-terminal region is very low, which shows a high degree of evolutionary divergence (S4F Fig). Although the I-loop of the SMAD4 binding domain, which has been implicated as an important structure for SMAD4 binding, is not very well conserved in Fuss and its homologues, we and others have detected an interaction between SMAD4 with Fuss and Skor2, respectively [11,14,18]. The repressive action of Ski/Sno proteins is generally exerted by the recruitment of a protein complex containing HDAC1 [10]. Skor1 and Skor2 also interact with HDAC1 and interestingly, it has been shown that the residues important for this interaction are localized in a segment reaching from amino acid 385-592 in mouse Skor2 [16,17]. Similar to the lack of the I-loop sequence, this segment is highly diverse between Fuss and Skor2 challenging if Fuss nevertheless is able to interact with Rpd3, the HDAC1 homologue in *Drosophila melanogaster* (S4F Fig). Therefore, we performed Co-Immunoprecipitations (CoIP) and transfected S2R+ cells with Fuss and Rpd3 tagged with FLAG or HA. Interaction between Fuss and Rpd3 could be shown independent of the type of the tags (Fig 6A). Skor1 and Skor2 have also been described to interact with Smad2 and Smad3, homologues of the *Drosophila* Smox, which executes the same function as Mad, but in the TGF-ß like signaling pathway [13,14,22]. Using the same methological approach as for the Fuss and Rpd3 interaction, we could not detect any interaction between Fuss and Smox, independent of the tags used (Fig 6A). Interestingly Smox is one of the genes specifically enriched in our TaDa datasets for Fuss neurons, so there would be a possibility for interaction in these cells.

**Fig 6.**
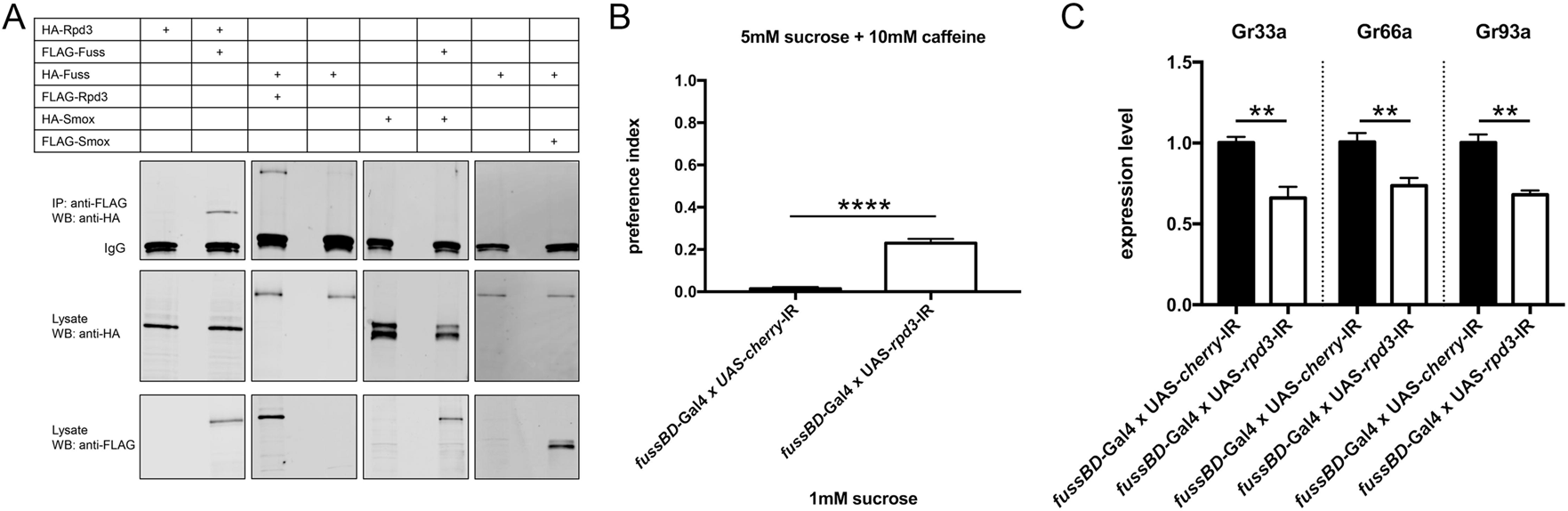
Rpd3 interacts with Fuss and phenocopies *fuss* mutant phenotypes. (A) Co-Immunoprecipitation experiments show that Fuss-HA binds to Rpd3-FLAG and Rpd3-HA interacts with Fuss-FLAG, respectively. No interaction between Smox and Fuss can be found regardless of the tags. (B) Knockdown of *rpd3* results in an increased preference index towards 5mM sucrose mixed with 10mM caffeine compared to *fussBD*-GAL4 x UAS-*cherry*-IR flies. n=4 for each genotype. One-way ANOVA with *post hoc* Tukey’s test was used to calculate p-values. ***p<0.001. ****p<0.001. Error bars indicate SEM. (C) Bitter gustatory receptors Gr33a, Gr66a and Gr93a are downregulated in *fussBD*-GAL4 x UAS-*rpd3*-IR flies compared to *fussBD*-GAL4 x UAS-*cherry*-IR flies. n=4-5 for each genotype. Unpaired t-test was used to calculate p-values. **p<0.01. Error bars indicate SEM.

If Fuss is acting within a protein complex in concert with Rpd3, we should be able to mimic *fuss* mutant phenotypes with *rpd3* depletion. Therefore, a UAS-*rpd3*-IR knockdown line was specifically expressed in Fuss neurons using the *fussBD*-Gal4 driver to reduce *rpd3* expression throughout development. Adult flies were then tested again in a two-choice feeding assay for bitter sensing. *Rpd3* knockdown flies showed a significant higher preference towards caffeine than control flies (*fussBD*-Gal4 x UAS-*cherry*-IR; Fig 6B). Because Rpd3 is involved in many different chromatin complexes, we analyzed again the expression levels of bitter gustatory receptors. Expression of all three tested GRs Gr33a, Gr66a and Gr93a was again diminished (Fig 6C) and therefore we conclude, that the Fuss/Rpd3 complex plays a key role in the final cell fate determination of gustatory neurons.

### Fuss function in CNS neurons – a contentious issue

In overexpression experiments, Ski/Sno proteins have often been identified as negative regulators of TGF-ß or BMP-signaling [14,17]. In *Drosophila*, Dpp is the main homologue to vertebrate BMPs and it is involved in multiple developmental signaling events, in particular in the *Drosophila* wing [49]. We have previously shown, that an overexpression of Fuss during wing development indeed results in diminished expression of Dpp target genes and, concomitantly, induces a phenotype, which resembles loss of Dpp signaling, despite the fact, that we could only detect a physical interaction with the Co-Smad Medea but not with the R-Smad Mad [18]. In Dpp signaling, Mad gets phosphorylated by the type I receptors Saxophon and/or Thick veins and, thus phosphorylated Mad is an excellent marker for active Dpp signaling and also for motoneurons or Tv neurons [50,51]. To analyse a possible role of Fuss in Dpp signaling, we used *fuss*^*Mi13731*^-flies, in which GFP is expressed under the *fuss* promotor to label Fuss expressing cells and we counterstained 3^rd^ instar larval brains with an antibody against phosphorylated Mad (pMad) (Fig 7A and Fig 7B). These results clearly showed that Fuss expression is not overlapping with pMad in heterozygous *fuss*^*Mi1373*^*/+* conditions. As there is a possibility that Fuss is acting upstream of Mad phosphorylation, we compared pMAD staining of heterozygous (Fig 7C) with homozygous *fuss*^*Mi13731*-^flies (Fig 7D). Again, there is no overlap of pMAD and GFP stainings in both genotypes, indicating that there is no increase of pMAD in *fuss* mutant neurons in the absence of Fuss. Importantly, this is in agreement with our overexpression studies, where Fuss had no influence on Mad phosphorylation [18]. Therefore, we conclude, that endogenously Fuss is not involved in Dpp signaling inhibition and it also emphasizes previous results, that Fuss is expressed in interneurons and not in motoneurons, which require pMad activity [51].

**Fig 7.**
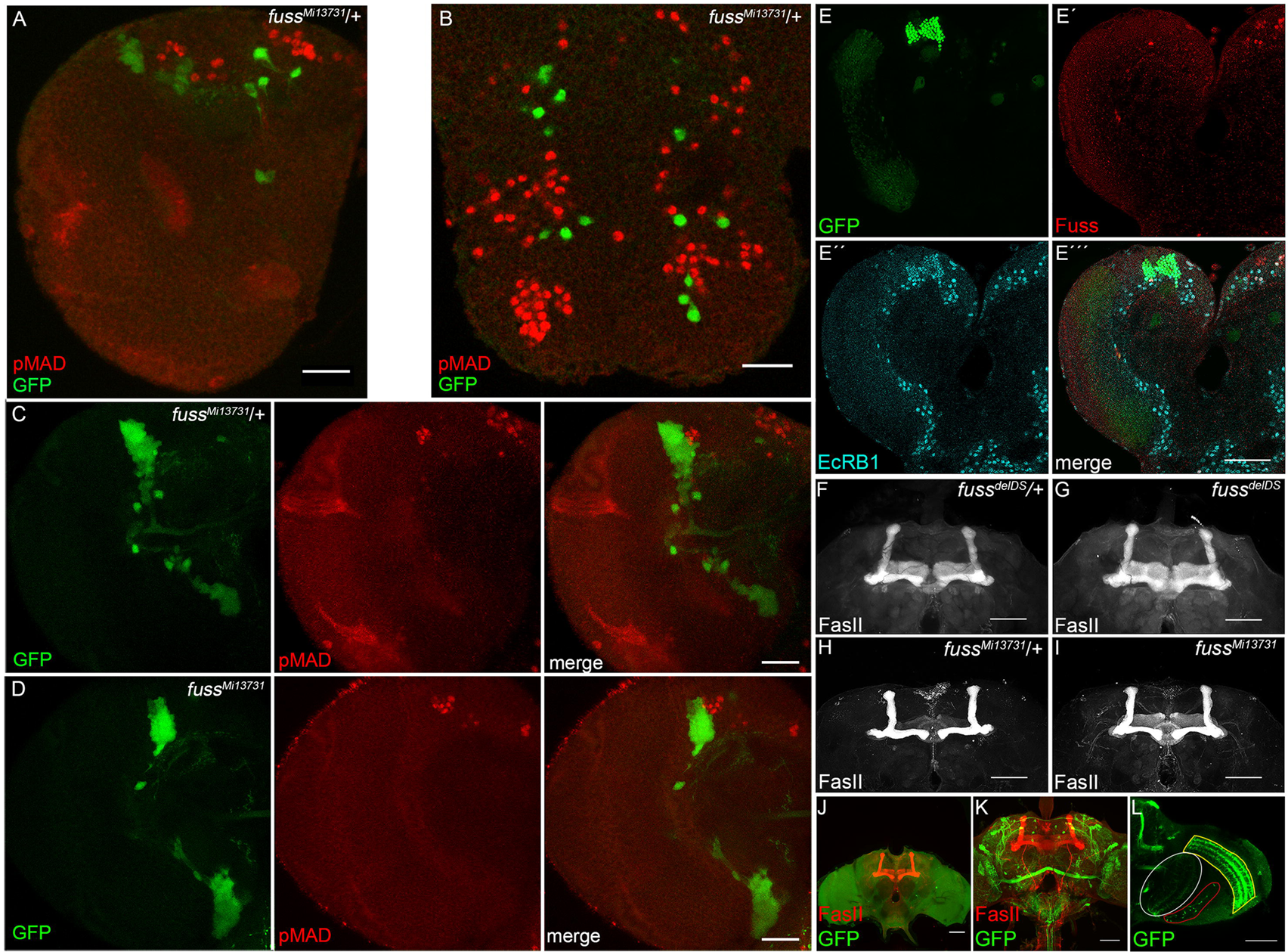
Fuss is neither a regulator of Dpp signalling nor involved in mushroom body formation. (A) GFP (green) expressed from heterozygous *fuss*^*Mi13731*^/+ reporter line does not colocalize with pMAD (red) in larval brain. (B) GFP (green) expressed from heterozygous *fuss*^*Mi13731*^/+ reporter line does not colocalize with pMAD (red) in larval VNC. (C) GFP (green) expressed from heterozygous *fuss*^*Mi13731*^/+ reporter line and homozygous *fuss*^*Mi13731*^ marks Fuss neurons. Anti-pMAD (red) displays active Dpp signaling. No colocalization can be observed in any genotype indicating, that Fuss itself is not involved in Dpp signaling inhibition. All pictures depict slices of the larval brain or VNC and not full stacks to exclude false positive colocalization. Scale bars indicate 25 *µ*m. (E) Representative picture of Kenyon cell nuclei of a third instar larval brain hemisphere. Nuclei of Kenyon cells are marked by colocalization of nuclear GFP driven by *OK107*-Gal4 (green, E) and EcRB1 (blue, E’’) in 3^rd^ instar larval brains. Anti-Fuss (red, E’) staining cannot be observed in the Kenyon cell clusters. Fuss cells positive for EcRB1 expression, do not overlap with GFP expression from *OK107*-Gal4 driver (E‴). (F-I) Mushroom bodies of an adult brain of heterozygous *fuss*^*delDS*^/+ flies (F), homozygous *fuss*^*delDS*^ flies (G), heterozygous *fuss*^*Mi13731*^/+ flies (H) and homozygous *fuss*^*Mi13731*^ flies (I) visualized by anti-FasII staining. (J) Ablation of Fuss neurons removes all Fuss neurons but mushroom body (red) stained with anti-Fas2 is not affected in adult brains. (K) *fussBD*-Gal4 driven UAS-*CD8-GFP* visualises projection pattern of Fuss neurons in an adult brain. (L) *fussBD*-Gal4 driven UAS-*CD8-GFP* shows Fuss neurons strongly project to lobula (white), lobula plate (red) and medulla (yellow) in an adult brain. Scale bars indicate 50 *µ*m.

Previously, the only loss of function data of *fuss* was generated using a genomic deletion of 40 kb including the *fuss* locus and additional genes [22]. This deletion lead to a reduced survivability during development, a shortened lifespan of the escapers and an impaired mushroom body development. All these phenotypes were attributed to the loss of Fuss expression. As we did not observe an impact on survivability or lifespan upon the loss of Fuss (see above), we wondered if Fuss is indeed involved in mushroom body development. Based on RNA *in situ* hybridisations Takaesu et al. assumed that Fuss is expressed in Kenyon cells during development and is required for the proper formation of the mushroom body [22]. Having now specific antibodies, gene trap constructs and *fuss* mutations in hand, we decided to carefully reevaluate this data on mushroom body expression and function during development. In a first step, we used *OK107*-Gal4 driven nuclear GFP as a marker for developing Kenyon cells and colabeled larval brains with EcRB1 and Fuss. We found that Fuss is not expressed in the developing mushroom body Kenyon cells, but it shows a partial overlap with EcRB1 expression outside of the Kenyon cell domain (Fig 7E-E‴). Next, we analysed adult mushroom body structures of *fuss* mutant flies using a FasII-antibody. As expected, due to the lack of Fuss expression in Kenyon cells, no deformation or loss of any of the lobes of the mushroom body was observed in homozygous *fuss*^*Mi13731*^ or *fuss*^*delDS*^-flies (Fig 7F–Fig 7I). In addition, the expression of *rpr* with the *fuss*^*delDS*^-Gal4 line lead to a complete ablation of Fuss neurons, but did not result in a malformation of adult mushroom bodies (Fig 7J). Furthermore, expression of CD8-GFP with *fussBD*-Gal4 in adult brains shows that Fuss neuron clusters are also localized distal to the mushroom body (Fig 7K). In fact, Fuss neuronal projections are localized outside of the mushroom body lobes in the adult brain and some Fuss neurons are targeting the optic lobe including different layers of the medulla, lobula and lobula plate but not the lamina (Fig 7L). From these results, we conclude that *fuss* has no impact on mushroom body development and that most of these neuronal populations such as the Fuss/Atonal positive neurons are higher order neurons of the visual system.

## Discussion

The molecular and cellular functions of the *fuss* genes, which are members of the Ski/Sno protein family, are still poorly understood. The fact that *Drosophila* contains only one single *fuss* gene offers a great opportunity for a thorough analysis. However, this has been restrained due to its location on the 4^th^ chromosome, where only limited genetic tools were available. As a consequence, previous reports have been focusing on the analysis of either overexpression studies or by using a multi-gene deficiency with contradictory results [18,22]. In the meantime, more recent methodological advances like the CRISPR/Cas9 genome editing [52] and the MiMIC gene trap technique [27] have expanded the *Drosophila* genetic toolbox and provided an appropriate genetic environment allowing a thorough and in-depth study of such genes. The availability of the *fuss*^*Mi13731*^ fly line, which is a gene trap of *fuss*, allowed us to study the expression pattern of Fuss. This line perfectly matches our Fuss-antibody stainings and was used to create a Gal4 line via RMCE as previously described [27]. A second independent mutant *fuss* allele, *fuss*^*delDS*^ was created by CRISPR/Cas9 editing by deletion of the main functional protein domains. Although *fuss*^*Mi13731*^ and *fuss*^*delDS*^ alleles are generated by different genetic approaches they share the same phenotypes, underlining that despite the complex genomic organization of *fuss* the observed phenotypes are due to the loss of *fuss*. Surprisingly, *fuss* mutant flies are fully viable and do neither show developmental lethality or reduced lifespans nor any other apparent phenotypes.

By means of our new tools, we could show that Fuss is expressed postmitotically in a small subset of neurons. All Fuss neurons in the CNS are interneurons, but they express different cell fate markers, suggesting that they represent a rather diverse group of neurons. These results were confirmed molecularly by a targeted DamID experiment, which, in addition, indicated a highly specific expression of gustatory receptor genes and indeed, Fuss is expressed in one GRN per sensillum. In S and I-type sensilla it is expressed in bitter GRNs and in L-type sensilla, which lack bitter GRNs, it is expressed in salt attracting GRNs. We investigated how the bitter GRNs react to the loss of Fuss and interestingly, this leads to an impairment of bitter sensation. Remarkably, this phenotype is correlated with a downregulation of bitter gustatory receptors Gr33a, Gr66a and Gr93a and in some bitter GRNs of *fuss* mutant flies no Gr33a expression can be observed anymore. The expression of Fuss in sensory neurons during development, and the adult phenotype, suggest that Fuss is needed for the proper maturation of these neurons and therefore is essential for bitter GRN differentiation. As there is a possibility, that the bitter sensation phenotype might be due to some higher order interneurons within the CNS, we generated a specific UAS-*t::gRNA-fuss*^*4x*^ line to be able to perform cell type specific gene knockouts. Indeed, using an independent driver line (*Poxn*-Gal4-13-1) expressed in all GRNs, faithfully reproduced this phenotype indicating a direct association of bitter sensation and GRN defects. In *fuss* mutant flies morphology of bitter GRNs was not altered and cell number was just slightly changed compared to controls, while Gr33a expression was completely lost in 40% of all bitter GRNs and Gr66a expression was reduced in all GRNs, but was never completely absent from a bitter GRN. Therefore, in *fuss* mutant flies bitter GRNs are correctly specified but the terminal differentiation of this neurons is disturbed, which ultimately results in impaired bitter taste sensation. This is comparable to Fuss neurons in the larval and adult CNS, where loss of Fuss expression also did not have an impact on axonal projections or cell numbers and thus not on initial specification of these neurons. This supports the idea, that Fuss is required for fine tuning individual subgroups of neurons during development, a phenotype, which resembles loss of Skor2 in mice, where it is dispensable for initial Purkinje cell fate specification but is required for proper differentiation and maturation of Purkinje cells [15]. It is very likely that other genes will also be affected by the loss of Fuss, and the reduction of these gustatory receptors could lead to a cumulative effect, as it has been shown that they act in heteromultimers where a multimeric receptor consists of at least Gr66a, Gr33a and Gr93a, which are all required for caffeine sensation [53,54]. Whereas over the years many studies have dissected the function of single gustatory receptors, the complexes they establish, and genes which are involved in more common topics like sensory neuron formation, less is known about the differentiation and specification of subsets of GRNs [55–57]. To find further genes involved in differentiation of bitter GRNs and to clarify the molecular consequences of the *fuss* mutation in bitter GRNs we will conduct transcriptional profiling experiments specifically in Fuss positive GRNs.

Using the TaDa method, we were curious to see if this method is sensitive enough to pick up significant differences between *fuss* mutant and wildtype flies. This was indeed the case for *GR66a*. However, in general, the performed TaDa experiments showed only slight differences between mutant and control flies. This could be a consequence of Fuss being expressed in heterogenic neuronal clusters. We showed, that Fuss interacts with Rpd3, a histone deacetylase, and therefore, a chromatin modifier, which is preferentially associated with inhibitory gene regulating complexes [58]. This could be a common mechanism for Fuss in all Fuss expressing neurons. However, different neuronal populations have different open and closed chromatin and probably the Fuss/Rpd3 complex regulates different genes in different neuronal populations, which could lead to the masking of differential gene expression by individual neuronal cell groups. Additionally, although the TaDa technique functions very well to generate transcriptional profiles without cell isolation, data is nondirectional and at GATC fragment resolution, which decreases overall resolution. To overcome these limitations experiments are on the way to unravel the function of specific neuronal clusters as well as the function of *fuss* in these neuronal clusters, and to specifically profile transcription of these clusters and changes upon loss of *fuss*.

A careful analysis with our newly generated antibodies shows, that there is no expression of Fuss in larval or adult Kenyon cells as has been postulated recently [22]. To unequivocally show, that there is no requirement for Fuss in mushroom body development, neither autonomously nor non-autonomously, Fuss expressing neurons were ablated using a *fuss*-GAL4 line driving Reaper. Again, these flies, even without any *fuss* expressing cells, are fully viable and do not show mushroom body defects. Lastly, we also did not find any evidence of Fuss being expressed in insulin producing neurons by our antibody staining or DamID experiments as shown recently [36]. These discrepancies are most likely explained by the use of the specific knockout line *fuss*^*delDS*^, and the gene trap line *fuss*^*Mi13731*^ in our case, whereas a 40 kb genomic deletion *Df(4)dCORL* was used in Takaesu et al. [22] and Tran et al. [36]. This deletion covered the *fuss* locus as well as two more protein coding genes, *4E-T* and *mGluR*, and three noncoding RNA genes, *CR45201*, *CR44030* and *sphinx*. Any of these, or a combination of them, could be responsible for premature lethality or mushroom body defects. One additional possible explanation for their mushroom body defects in the deletion is an inappropriate fusion of a new transcriptional start site or enhancer region from the *mGluR* upstream to the *toy* gene creating a weak overexpression phenotype of toy in mushroom bodies, a phenotype, which has been described already [59]. Indeed, very recently Tran et al. [35] described a slight overexpression of Toy in their deficiency allele *Df(4)dCORL*.

We and others have shown that Ski/Sno protooncogenes have an inhibitory effect on TGF-ß or BMP signaling in overexpression assays [18,60]. This is often associated with the ability of Ski/Sno proteins to inhibit the antiproliferative effects of TGF-ß signaling in cancer and to promote their progression [61]. However, in an endogenous situation, Fuss is not expressed in cells, where the BMP/Dpp signaling pathway is active. This is displayed by the absence of the motoneuron marker pMad in Fuss neurons. Later in adulthood, Mad itself is also not specifically enriched in Fuss expressing neurons according to the TaDa dataset, clearly pointing against a function in BMP signalling. We also tested if Fuss is involved in the Activin signaling cascade, but we could not detect an interaction between Fuss and Smox in CoIP assays. However, we cannot rule out the possibility that the phosphorylated form of Smox is interacting with Fuss or the Fuss/Med complex. But since both, phosphorylated Smox and Fuss interact with Medea, we would potentially also get an artificial interaction [18,62]. At least according to the TaDa dataset, Smox is expressed in Fuss neurons. Unfortunately, there is currently no good marker available to test for an activated TGF-ß signaling pathway in *Drosophila* cells, like an antibody against phosphorylated Smox. What might be the main molecular mechanism for Fuss? Although the Ski/Sno/Dac homology domain and the SMAD4 binding domain in Ski have DNA binding character, they mainly have been shown to be involved in protein-protein interactions [11,63]. Furthermore, Ski/Sno proteins do not possess an intrinsic catalytic activity, they rather act as recruiting proteins [2]. In agreement, we could show that this is also the case for Fuss. Not only that Fuss binds to Medea, which is a DNA binding protein and therefore mediates the DNA binding, Fuss also interacts with Rpd3, a histone deacetylase. Thus, the Med/Fuss/Rpd3 complex is involved in chromatin silencing and plays a key role in terminal differentiation. Interestingly, the loss of bitter sensation and downregulation of bitter GRs could also be phenocopied by a knockdown of *rpd3* in Fuss expressing gustatory neurons. One current hypothesis of Fuss/Rpd3 function in GRNs, which we propose, is, that this protein complex is inhibiting a repressor of GR genes and in the absence of either *fuss* or *rpd3*, the complex is inactivated, this repressor will inhibit bitter GR genes.

For Ski and Sno, the transcriptional repressor complexes have been reasonably well characterized [10,64], but for the Fuss-type proteins, very little is known about their complexes. It would be highly interesting if Fuss proteins act through repressor complexes identical to the complexes of Ski or Sno or a rather unique one. The most exciting question to solve regarding protein interaction will be, if the Fuss/Rpd3 complex plays a role in TGF-ß signalling, or if in contrast to its mammalian homologues, it is not only acting BMP independent, but also independent from the TGF-ß signalling cascade. Besides identifying further protein-protein interactions and investigating DNA-protein interactions more precisely, it will be very important to describe the exact function of the Fuss/Rpd3 complex. In mammals, Skor2 is thought to activate Sonic Hedgehog expression in Purkinje cells from direct binding to the Sonic Hedgehog promotor and this might be achieved by inhibition of the BMP pathway or by cooperation with the RORalpha pathway, a nuclear orphan receptor [15,17]. In contrast to that, Skor1 interacts with Lbx1, a homologue of the ladybird early or ladybird late in *Drosophila*, and acts as a transcriptional corepressor of Lbx1 target genes [16]. Our TaDa datasets strongly point towards another function for Fuss in *Drosophila*, as neither *hedgehog* nor the homologues of *Lbx1*, *ladybird late* and *ladybird early*, are enriched in Fuss expressing cells. Therefore, identifying target genes, interacting proteins, binding motifs of the Fuss complex and subsequent comparison with established models for other transcription factor complexes will elucidate the role of this complex in cell fate determination.

## Material and Methods

### *Drosophila* genetics

Flies were kept under standard conditions (25 °C, 12 h/12 h LD cycle). Flies from RNA interference crosses were kept at 29 °C. Fly lines obtained from the Bloomington Stock Center were *fuss*^*Mi13731*^ (#60860), UAS-*CD8-GFP* (#5137), UAS-*CD8-RFP* (#32218), UAS-*LacZ* (#8529), *tubulin-*Gal80ts (#7108), UAS-*Stinger* (#65402), UAS-*rpd3*-IR (#33725), UAS-*cherry*-IR (#35785), *ap*-Gal4 (#3041), *Gr33a*-Gal4(#31425), *Gr66a*-Gal4 (#57670), *Gr93a*-Gal4 (#57679), *Hb9*-Gal4 (#32555), *Gr5a*-Gal4 (#57591), *Ir76b*-Gal4 (#51311), UAS-*cas9* (#58985) and *ato*-Gal4 (#6480). UAS-*Dam* and UAS-*Dam-PolII* stocks were a gift from Andrea Brand. *Poxn*-Gal4-13-1 was a gift from Markus Noll. UAS-*fussB*, *ap*-tau-LacZ and UAS-*rpr* were from our stock collection. To generate the *fuss*^*delDS*^ line two sgRNAs (GTAAGCTCCGTTTTGCTGTA and GGTGTTCCCTTTAACTTACA) were employed and cloned into *pU6-BbsI-chiRNA*. Homology arms were cloned into *pHD-DsRed-attP* and coinjected with *pU6-BbsI-chiRNA* as described in Gratz et al. [52]. The *fussBD*-Gal4 and the *fuss*^*Mi-cherry*^ lines were created via RMCE with the vectors *pBS-KS-attB1-2-GT-SA-GAL4-Hsp70pA* and *pBS-KS-attB1-2-GT-SA-mCherry-SV40, respectively* [27]. To generate the mutant *fuss*^*delDS*^-Gal4 line, the *fussBD*-Gal4 line was additionally targeted with the same sgRNAs via CRISPR/Cas9, which were used for the *fuss*^*delDS*^ line. Genomic DNA of CantonS and *fuss*^*delDS*^ flies was extracted with QIAamp DNA Mini Kit (Qiagen, Hilden, Germany). Successful indel mutation was confirmed by PCR with Cr1seqfw (CAAATCGACTGGGTAAATGGT) and Cr2seqrv (GTAGTCCACTACAAAGTTCCTG) oligonucleotides und subsequently sequenced (GATC Biotech, Konstanz, Germany). *hs*-*fussB-GFP* was generated by cloning the ORF of *fussB-GFP* into *pCaSpeR. hs*-*fussB-GFP* flies were generated via P-element integration of *pCaSpeR-hs-fussB-GFP* vector into *w; +/Δ2-3, Ki* and subsequent crossed to W^1118^ flies and transformants were balanced. For generation of UAS-*t::gRNA-fuss*^*4x*^ flies we followed the protocol from Port et al. [28] and used primers, which allow the targeting of the CRISPR target sites GTAAGCTCCGTTTTGCTGTACGG, ATTGTATCCCTGCACATTGAAGG, CCAGTGAGTTCCCGACGATGTGG and TTGAAATTTGCGCCAAGCAAAGG. The *pCFD6*-*t::gRNA-fuss*^*4x*^ was injected into *y[1],M{vas-int.Dm}ZH-2A,w[*]; M{3xP3-RFP.attP}ZH-86Fb* flies to generate UAS-*t::gRNA-fuss*^*4x*^ flies. All *Drosophila* strains generated in this publication are available upon request.

### Polyclonal anti-Fuss antibody generation

Fulllength *fuss* ORF was codon optimized at GeneArt, Regensburg, Germany. An appropriate fragment of the codon optimized fuss gene was cloned into pQE60 resulting in a 16 kDa 6xHis tagged Fuss fragment called Fuss16-6xHis (Fig S1). Transformed Rosetta2 cells were grown to an OD 0.6 and protein expression was induced with 0.5 mM IPTG. Cells were incubated for 2.5 h at 37 °C, harvested, resuspended in PBS supplemented with Protein Inhibitors (Roche, Switzerland) and lysed via sonication. Fuss16-6xHis was purified with an Äktapurifier10 (GE Healthcare, Life sciences) and was used for immunization of two rabbits at Davids Biotechnologie, Regensburg, Germany. The resulting antiserum was purified against Fuss16-6xHis to reduce nonspecific binding. Before using the anti-Fuss antibodies for immunostainings or western blots they were preabsorbed using 0-6 h embryos treated with 4 % PFA in PBST 0.1 % as follows: The antibody was diluted to 1:50 in 500ml PBST 0.1 %, NGS 5 % and incubated with 100 *µ*l fixed embryos on a rotator at 4 °C over night. Anti-Fuss antibody was further diluted to 1:200 in PBST 0.1 %, NGS 5 % for immunostainings and 1:1000 in TBST 0.1 % for western blots.

### Real time PCR

Sixty proboscises from each genotype (equal number of males and females) per biological replicate were dissected on ice and snap-frozen in liquid nitrogen. RNA was extracted by adding lysis buffer from the MicroSpin Total RNA Kit (VWR) and the tissue was extracted with a bead mill and it was proceeded according to the manufacturer’s protocol. cDNA was generated with the QuantiTect^®^ Reverse Transcription Kit (QIAGEN). For subsequent real time PCR ORA qPCR Green ROX L Mix (HighQu, Kralchtal, Germany) was employed. RP49 was used as a housekeeper control, with the primers RP49fw (CCAAGCACTTCATCCGCCACC) and RP49rv (GCGGGTGCGCTTGTTCGATCC). Primer sequences for Gr33afw (CCACCATCGCGGAAAATAC), Gr33arv (ACACACTGTGGTCCAAACTC), Gr66afw (ACAGGAATCAGTCTGCACAA), Gr66arv (AATGTTTCCATGTCCAGGGT), Gr93afw (CCACGTCACAAACTCATTCC), Gr93rv (GCCATCACAATGGACACAAA), fussBDfw (TGGCTTCTATATCTGTGGCTCA) and fussBDrv (CAAAGGCGCTCTTGACCTTC) were generated with PrimerBlast. For relative quantification, we applied the ΔΔCT method. Every experiment has been repeated at least four times.

### Protein expression analysis

Developmental studies Hybridoma Bank (DSHB) antibodies were: Acj6 (1:50), Dac (Mabdac1-1, 1:20), EcRB1 (AD4.4, 1:50), LacZ (JIE7, 1:20), Pros (MR1A, 1:10), Elav (7E8A10, 1:50), Engrailed (4D9, 1:20), Even skipped (3C10, 1:20) Repo (8D12, 1:20), and Fas2 (1D4 1:10). Additional antibodies were: Pale (AB152, 1:500, Millipore), Ilp2 (1:400, gift from Pierre Leopold), Toy (1:200, gift from U. Walldorf), GFP (goat 1:100, Rockland; rabbit 1:1000, ThermoFisher), RFP (rabbit 1:20, ThermoFisher) and anti-phospho-SMAD1/5 (1:50, Cell signaling). Secondary antibodies were goat anti-mouse, anti-rabbit, anti-rat and anti-guinea pig Alexa Fluor 488, 555 and 594 (ThermoFisher). Samples were analysed with a Leica SP8 microscope. To confirm functionality of anti-Fuss antibodies *hs*-*fussB-GFP* third instar larvae were heatshocked for one hour at 37 °C and were allowed to recover for another hour at room temperature. RIPA buffer was added to ten larvae and they were mechanically disrupted. Insoluble fragments were removed by centrifugation and supernatant was incubated at 95 °C for five minutes. Supernatant was analysed via SDS-Page and Western blotting. As a housekeeper mouse anti-tubulin (B-5-1-2, MERCK) was utilised and secondary antibodies were goat anti-mouse 680nm and goat anti-rabbit 800nm (Li-Cor, Lincoln, USA). Signals were detected using an Odyssey infrared imaging system (Li-Cor, Lincoln, USA).

### Co-Immunoprecipitation

S2R+ cells were cultured in Schneider’s *Drosophila* Medium (Pan Biotech, Aidenbach, Germany) supplemented with 10 % Fetal Bovine Serum (Pan Biotech, Aidenbach, Germany). The coding regions of *fussB, smox* and *rpd3* were inserted into pFSR11.58 3xHA and pFSR12.51 4xFlag (Frank Sprenger, Regensburg, Germany). Cells were transfected in 6 well plates at 70 % confluency with 2 *µ*g of pFSR11.58 Fuss-HA and pFSR12.51 Rpd3-Flag (or Smox-Flag), or pFSR11.58 RPD3-HA (or Smox-HA) and pFSR12.51 Fuss-Flag, respectively, using Lipofectamine 3000 (Thermo Scientific, Waltham, MA, USA) according to the manufacturer’s protocol and incubated for another 24 h. Transfected cells were harvested using a plastic scraper. For Rpd3 and Fuss interaction experiments nuclear extracts were prepared using the NE-PER Nuclear and Cytoplasmic Extraction Reagents (Thermo Scientific, Waltham, MA, USA) and only nuclear fraction was used. For Fuss and Smox interaction whole cell extracts were prepared with 400 *µ*l lysis buffer (50 mM HEPES pH 7.5, 150 mM NaCl 150, 1 % Triton X-100, 10% Glycerol, 1 mM EGTA, 10 mM NaF) supplemented with cOmplete™ Mini Protease Inhibitor Cocktail (Roche, Switzerland). After preclearing the extracts with 30 *µ*l Protein A-Agarose beads (Santa Cruz, Dallas, TX, USA) and conjugating 1.5 *µ*l Anti-Flag M2 antibody (Sigma, St. Luis, Mo, USA) to 30 *µ*l Protein A/G Plus beads (Santa Cruz, Dallas, TX, USA), the volume of the nuclear extract was brought up to 400 *µ*l using RIPA buffer (50 mM Tris (pH 7.5), 150 mM NaCl, 1 % (v/v) NP-40, 0.5 % (w/v) Deoxycholat) supplemented with cOmplete™ Mini Protease Inhibitor Cocktail (Roche, Switzerland). 5% of the precleared extracts were saved for input analysis. Immunoprecipitation was conducted for 2 h at 4 °C. Following three washing steps with RIPA buffer, the precipitated proteins, as well as the precleared nuclear extracts, were analyzed by SDS-PAGE and western blotting. As primary antibodies Anti-Flag M2 and Anti-HA.11 (Covance Inc. USA) were used. Secondary antibody was goat anti mouse 680 (Li-Cor). Signals were detected using an Odyssey infrared imaging system (Li-Cor, Lincoln, USA).

### Targeted DamID and Bioinformatics

Targeted DamID to profile transcription in Fuss expressing neurons was performed as previously described [34,65,66]. UAS*-Dam,* UAS*-DamPolII,* UAS*-Dam; fuss*^*delDS*^ *or* UAS*-DamPolII; fuss*^*delDS*^ flies were crossed to *tubulin-*Gal80^ts^*; fuss*^*delDS*^*-*Gal4 flies. Three biological replicates of *DamPolII* expressing flies and three biological replicates of Dam expressing flies were conducted. Per replicate 100 one to three-day old flies (50 females and 50 males) were incubated for 24 h at 29 °C and snap-frozen in liquid nitrogen. Heads were detached by vortexing and separated with sieves. Processing of genomic DNA from heads and data analysis were performed as described and NGS libraries libraries were prepared with NEBNext UltraII DNA Library Prep Kit for Illumina [34,65,66]. Sequencing was carried out by the Biomedical Sequencing Facility at CeMM. For aligning reads, dm6 release from UCSC was used. Data tracks from same genotype were averaged with the average_tracks script and 3150 genes were called with an FDR < 0.01 for mutant flies and 2932 genes for control flies. log2(Dam-PolII/Dam) ratio datasets were visualized with the Integrative Genomic Browser.

### Life span

For life span determination, male flies were collected within 24 h after eclosion and were raised at 25 °C under a 12 h:12 h light/dark cycle. These flies were transferred to fresh food vials every two to three days.

### Two choice feeding assay

Feeding behaviour was analysed as previously described at 25 °C [38]. Fly age at time of testing ranged from one to three days and experiments were only accounted if at least 30 % of all flies showed clear evaluable coloured abdomen. As bitter compounds caffeine and denatonium benzoate were utilised at the indicated concentrations. Because feeding behaviour was influenced by temperature, *fussBD*-Gal4 x UAS-*cherry*-IR and *fussBD*-Gal4 x UAS-*rpd3*-IR flies were shifted to 25°C two hours prior testing. Every experiment has been repeated at least four times.

### Preparation of Figures

All figures were assembled with Adobe Photoshop CC (Adobe Systems) by importing microscopy images from Fiji and graphs from Prism.

### Statistics

Survival data were analyzed using the Log-rank (Mantel-Cox) and Gehan-Breslow-Wilcoxon tests. Significance was determined by two-tailed t-test or by One-way ANOVA with *post hoc* Tukey Multiple Comparison Test (****p<0.001; ***p<0.001; **p<0.01 and *p<0.05). Statistical analysis was carried out using Prism version 7.0a for MacOs, GraphPad Software, La Jolla, CA, USA.

### Availability

Raw sequencing data are accessible via Gene Expression Omnibus: GEO Series GSE115347.

## Supporting information

## Acknowledgements

We thank Juan A. Navarro for critically reading the manuscript. We thank Renate Reng for technical assistance. We thank Lars Kullmann as well as Caroline Iglhaut for the generation of *HS-FussB-GFP* flies and Matthias Schramm for the generation of the appropriate vectors for the *fuss*^*delDS*^ flies. We thank Frank Sprenger for providing pFSR vectors, Markus Noll for *Poxn-*Gal4-13-1 flies and Andrea H. Brand for UAS-*Dam* and UAS-*Dam*-*PolII* vectors and flies. We thank Michael Rehli and Owen J. Marshall for support regarding NGS library preparation and bioinformatical analysis. We would like to thank the Biomedical Sequencing Facility at CeMM for sequencing. Stocks obtained from the Bloomington Drosophila Stock Center (NIH P40OD018537) and antibodies from Developmental Studies Hybridoma Bank were used in this study.

## Supporting information

**S1 Fig. Characterization of *fuss*^*delDS*^ and *fuss*^*Mi13731*^ mutant flies.** (A) Genotyping of CantonS and homozygous *fuss*^*delDS*^ flies with fuss crispr1 seq fw and fuss crispr2 seq rv oligonucleotides via PCR of genomic DNA shows reduction of around 700bp in *fuss*^*delDS*^ mutants as expected in contrast to control (B). Staining of heterozygous *fuss*^*delDS*^/+ embryos with anti-Fuss (red) and anti-Elav (green) antibodies. Scale bar indicates 25 *µ*m. (C,D) Staining of homozygous *fuss*^*delDS*^ embryos with anti-Fuss (red) and anti-Elav (green) antibodies. Scale bar indicates 25 *µ*m. (B) Analysis of *fussB* and *fussD* transcript levels with fussBD fw and fussBD rv oligonucleotides via qPCR reveals a reduction of *fussB* and *fussD* transcript levels to 10 % in homozygous *fuss*^*Mi13731*^ flies in contrast to WTB flies. n=4 for each genotype. One-way ANOVA with *post hoc* Tukey’s test was used to calculate p-values. ****p<0.0001. **p<0.01. Error bars indicate SEM. (E) Anti-Fuss staining colocalizes with GFP in larval brains of heterozygous *fuss*^*Mi13731*^/+ line. (F) No anti-Fuss staining in larval brains of homozygous *fuss*^*Mi13731*^ line can be detected. Arrowhead indicates magnified cell cluster. (G) Location of the four CRISPR target sites of the UAS-*t::gRNA-fuss*^*4x*^construct in the DNA sequence of the Ski/Sno homology domain. (H) Adult brains of UAS-*cas9*/UAS-*fussB-GFP*; *fussBD*-Gal4/+ flies show normal Fuss expression pattern-Scale bar indicates 50 *µ*m. (I) In flies of the genotype UAS-*cas9*/UAS-*fussB-GFP*; UAS-*t::gRNA-fuss*^*4x*^*; fussBD*-Gal4/+ fussB-GFP is strongly reduced. Scale bar indicates 50 *µ*m.

**S2 Fig. Characterization of anti-Fuss antibody and Fuss neurons.** (A) Schematic representation of conserved domains and localization of the Fuss16 fragment used for immunization. Exact sequence of Fuss16-His fragment shown in red. (B) Detection of Fuss-GFP (green) from heatshock induced Fuss-GFP flies in western blots by anti-GFP and anti-Fuss antibodies. As a negative control CantonS is used and Tubulin as a housekeeper protein (red). Both antibodies recognize a predicted protein size of 112 kDa for the fusion protein. Endogenous levels of the Fuss protein cannot be detected on western blots due to the low abundance of the protein. (C) Comparison of VNC of stage 13 embryo with VNC of stage 16 embryo shows increase in number of Fuss (green) or Toy (red) cells, but only one cell per hemineuromer shows colocalization of both markers. (D) Comparison of expression pattern of interneuron marker Engrailed (red) and Fuss (green) visualized by antibody staining in embryonic VNC. (E) Fuss expression pattern as revealed by expression of GFP (green) by heterozygous *fuss*^*Mi13731*^/+ in larval brains does not colocalize with LacZ (red) driven by *Hb9*-GAL4 line. (F) Even skipped (red), a motor neuron marker, is not expressed in Fuss neurons (green) visualized by antibody staining in embryonic VNC. (G) Ventral Apterous cells marked by expression of CD8-RFP (red) with *ap*-Gal4 are positive for Fuss (green) expression in larval VNC. Scale bars indicate 30 *µ*m (C, D, E, F) and 10 *µ*m (G).

**S3 Fig. Fuss is not expressed in adult insulin like producing cells or dopaminergic neurons as revealed by TaDa and immunostainings.** (A) *pale* (*ple*) is weakly bound by Dam-PolII as revealed by TaDa and no colocalization is observed between Ple positive cells (red) and GFP expressed by the heterozygous *fuss*^*Mi13731*^/+ reporter line (green) in whole adult brains. Overlap between signals arises from different optical slices and not from colocalization. (B) *insulin like peptide 2* (*ilp2)* is weakly bound by Dam-PolII as revealed by TaDa. Confocal slices covering the pars intercerebralis and a part of the adult brain hemisphere show no colocalization between insulin producing cells labeled with anti-Ilp2 antibody (red) and Fuss neurons labeled with anti-Fuss antibody (green). In (A) and (B) regions bound stronger by Dam-PolII than by Dam are depicted in green, whereas regions bound stronger by Dam than by Dam-PolII are depicted in red. Scale bars indicate 50 *µ*m.

**S4 Fig. *fuss* mutant flies show an impaired bitter taste sensation.** (A) Expression of UAS-*CD8-GFP* with *fussBD*-Gal4 reveals four GRNs located in the two last tarsal segments of the prothoracic, mesothoracic and metatoracic leg. Scale bars indicate 50 *µ*m. (B) GFP expression from *Poxn*-Gal4-13-1 is not overlapping with Cherry expression from *fuss*^*Mi-cherry*^ reporter line in neurons of the adult CNS. Overlap can only be observed in GRN nerve fibers from proboscis. EBN = ellipsoid body neurons. ALI = Antennal lobe interneurons. Scale bar indicates 50 *µ*m. (C) Homozygous *fuss*^*Mi13731*^ flies show reduced caffeine sensation also at lower concentrations compared to heterozygous *fuss*^*Mi13731*^ x WTB and WTB flies. n=4-9 for each genotype. One-way ANOVA with *post hoc* Tukey’s test was used to calculate p-values. **p<0.01 ****p<0.0001. Error bars indicate SEM. (D) Homozygous *fuss*^*Mi13731*^ mutant flies show reduced sensation of denatonium benzoate compared to heterozygous *fuss*^*Mi13731*^ x WTB and WTB flies at a concentration of 100 *µ*m. At 500 *µ*m denatonium benzoate effect of homozygous *fuss*^*Mi13731*^ flies is reversed to control levels. n=4-5 for each genotype. One-way ANOVA with *post hoc* Tukey’s test was used to calculate p-values. ****p<0.0001. Error bars indicate SEM. (E) Transheterozygous *fuss*^*Mi13731*^/*fuss*^*delDS*^ mutants show reduced transcript levels fo Gr33a, Gr66a and Gr93a in contrast to W^1118^ control. n=4 for each genotype. One-way ANOVA with *post hoc* Tukey’s test was used to calculate p-values. ***p<0.001. **p<0.01. *p<0.05. Error bars indicate SEM. (F) Alignment of *Drosophila* Fuss with mouse Skor1 and Skor2. Ski/Sno/Dac homology domain, SMAD4 binding domain and proposed Rpd3 interaction fragment in Skor2 are displayed by colored lines as described.

**S1 Appendix. Average PolII occupancy and FDR of control dataset.**

**S2 Appendix. Average PolII occupancy and FDR of mutant dataset.**

**S3 Appendix. Data for generating graphs.**

