## Supplementary figures and images for "The *Drosophila fussel* gene is required for bitter gustatory neuron differentiation acting within an Rpd3 dependent chromatin modifying complex"

### Supplementary file 1

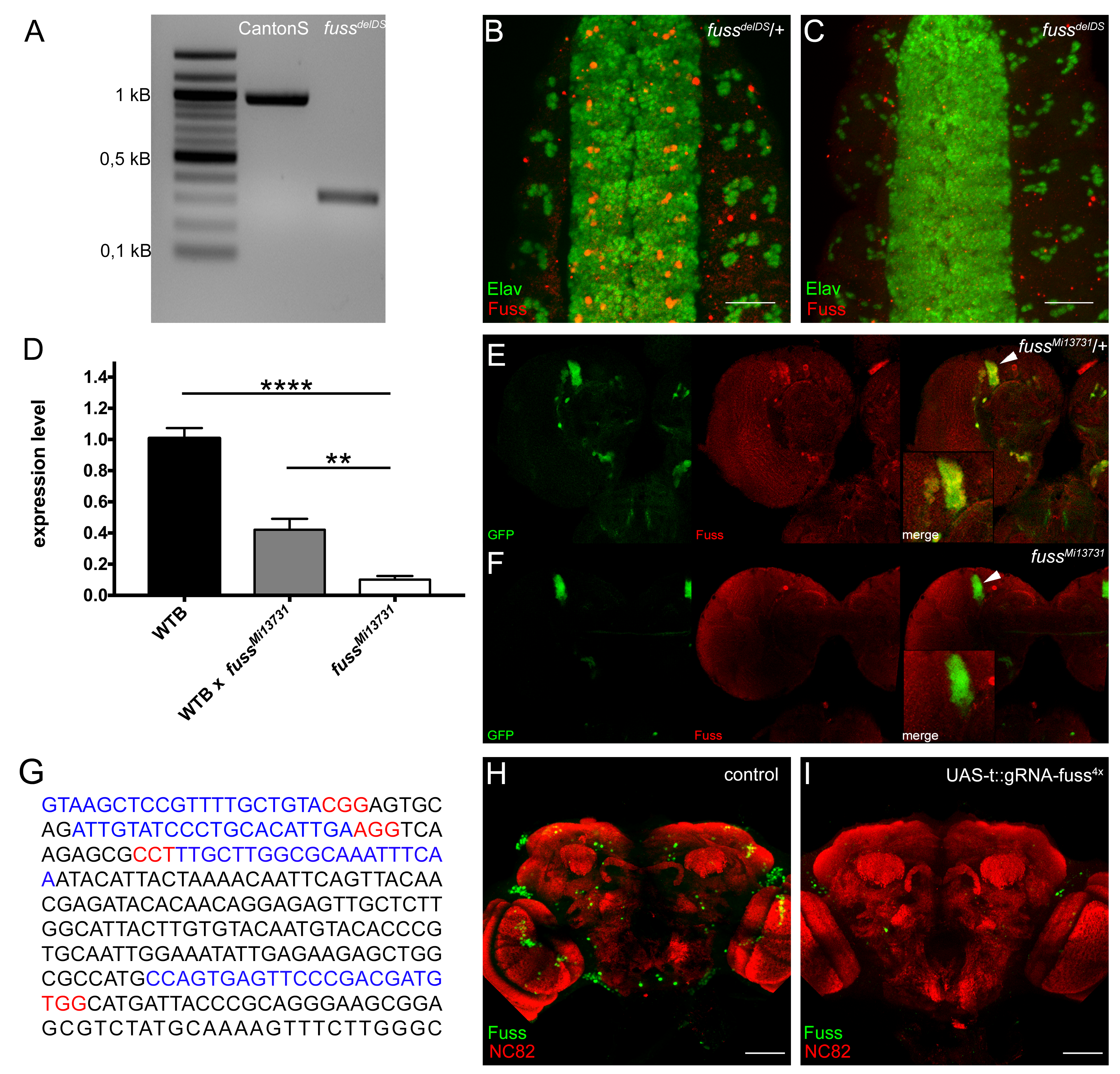

### Supplementary file 2

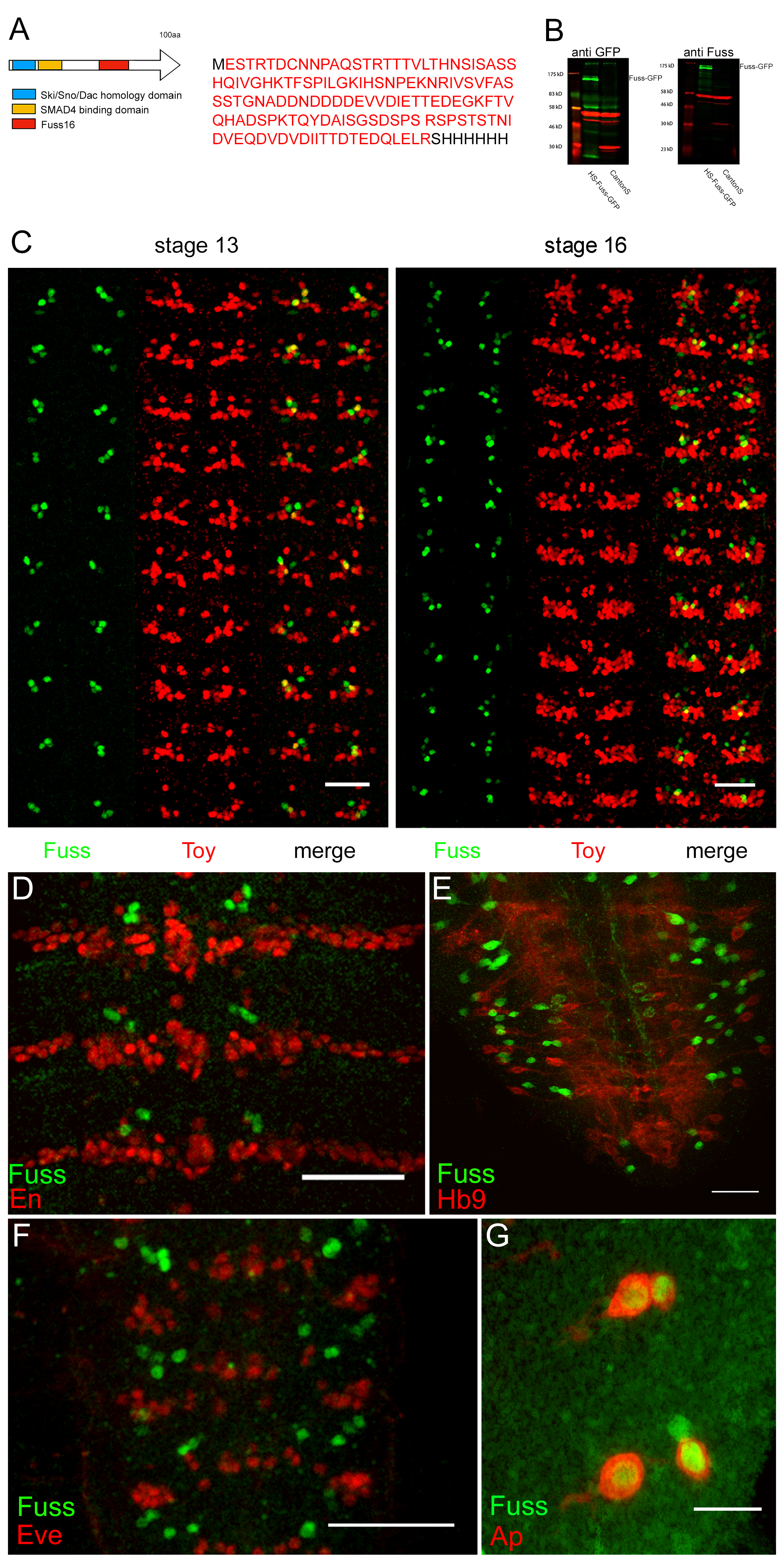

### Supplementary file 3

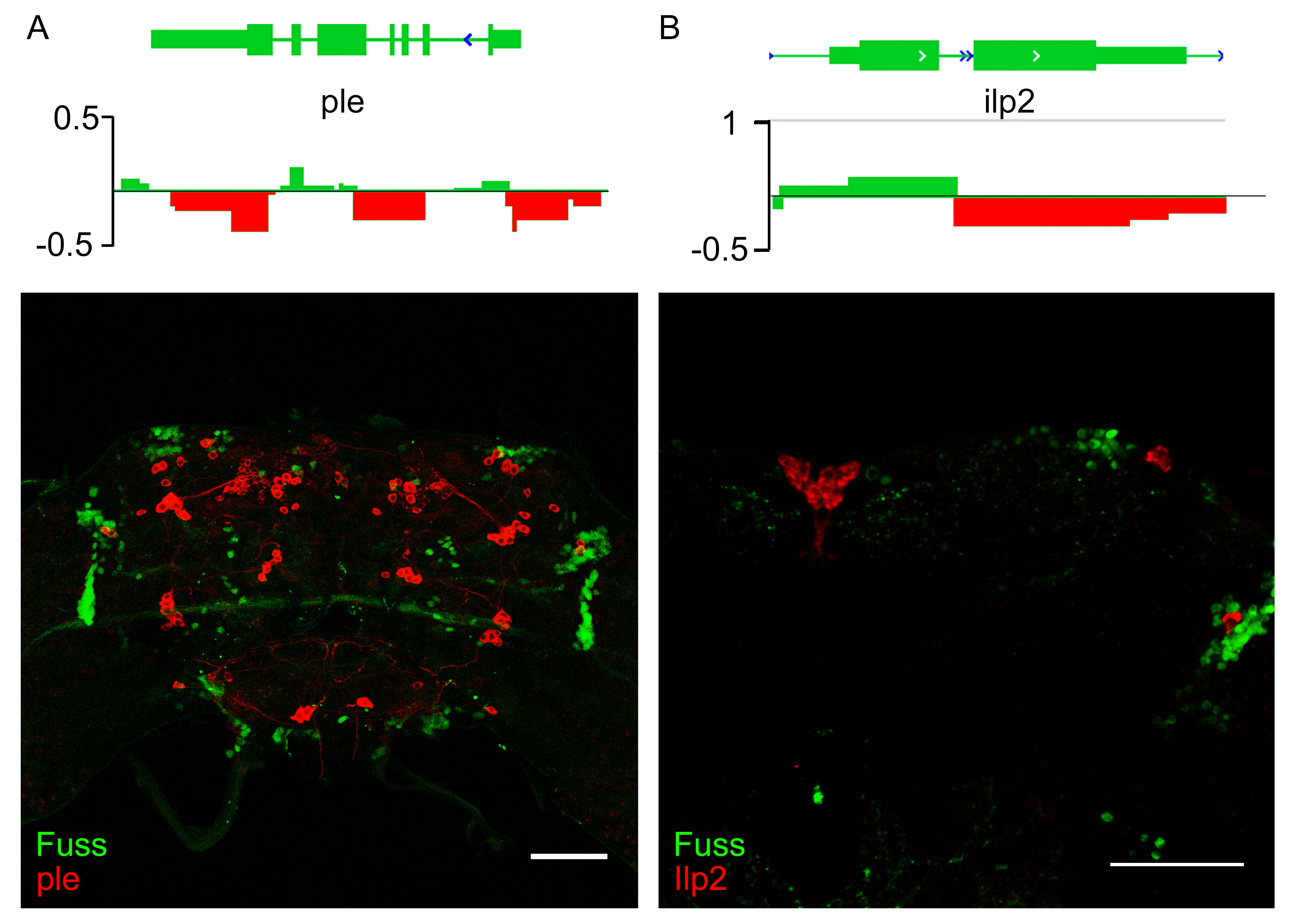

### Supplementary file 4

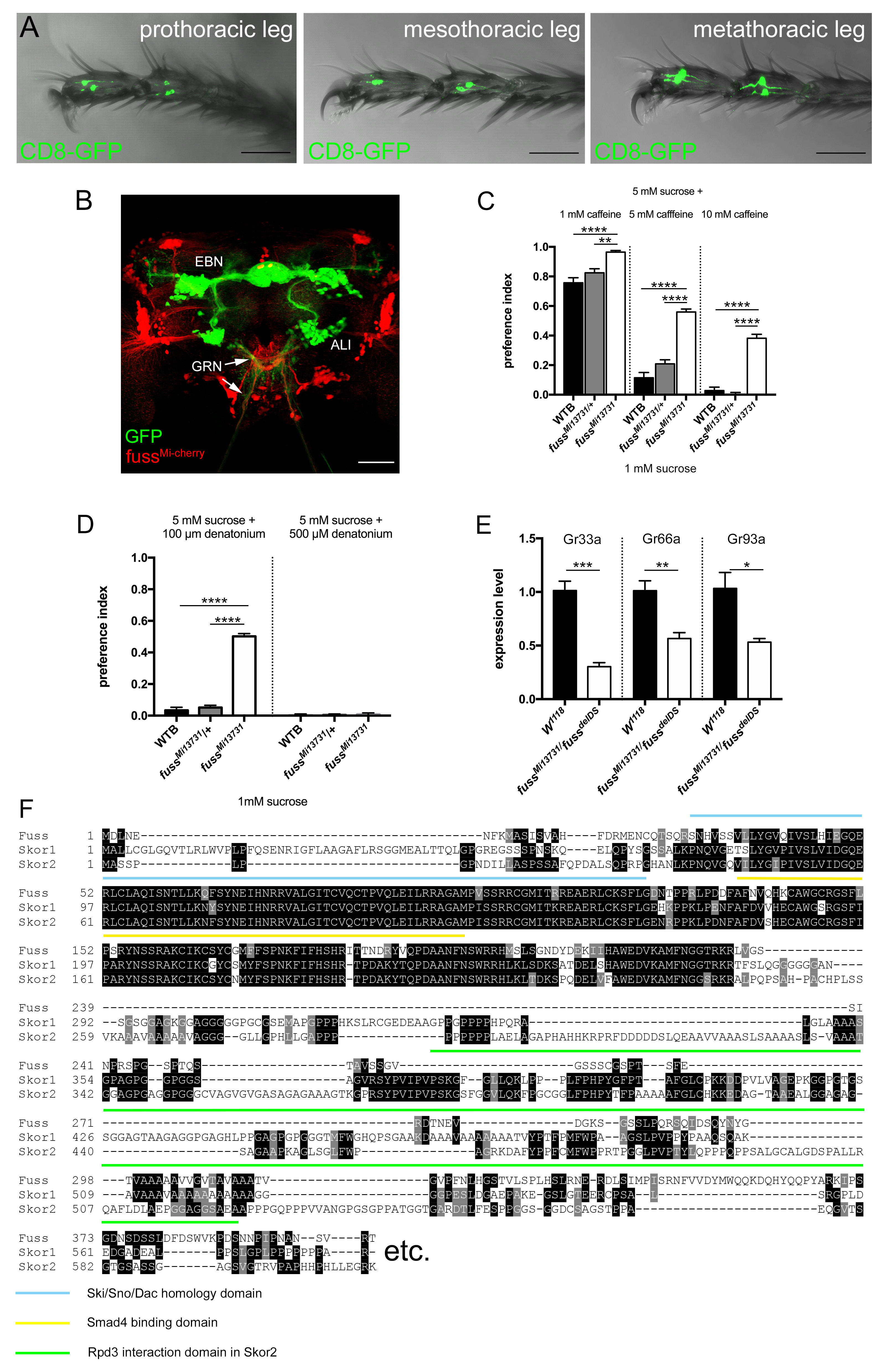
